# Fixation times of *de novo* and standing beneficial variants in subdivided populations

**DOI:** 10.1101/2023.07.07.548167

**Authors:** Vitor Sudbrack, Charles Mullon

## Abstract

The rate at which beneficial alleles fix in a population depends on the probability of and time to fixation of such alleles. Both of these quantities can be significantly impacted by population subdivision and limited gene flow. Here, we investigate how limited dispersal influences the rate of fixation of beneficial *de novo* mutations, as well as fixation time from standing genetic variation. We investigate this for a population structured according to the island model of dispersal allowing us to use the diffusion approximation, which we complement with simulations. We find that fixation may take on average fewer generations under limited dispersal than under panmixia when selection is moderate. This is especially the case if adaptation occurs from *de novo* recessive mutations, and dispersal is not too limited (such that approximately *F*_ST_ < 0.2). The reason is that mildly limited dispersal leads to only a moderate increase in effective population size (which slows down fixation), but is sufficient to cause a relative excess of homozygosity due to inbreeding, thereby exposing rare recessive alleles to selection (which accelerates fixation). We also explore the effect of meta-population dynamics through local extinction followed by recolonization, finding that such dynamics always accelerate fixation from standing genetic variation, while *de novo* mutations show faster fixation interspersed with longer waiting times. Finally, we discuss the implications of our results for the detection of sweeps, suggesting that limited dispersal mitigates the expected differences between the genetic signatures of sweeps involving recessive and dominant alleles.

## 1 Introduction

Populations can adapt to their environment via the fixation of beneficial alleles (Kimura and Ohta, 1971; Gillespie, 1994). Understanding the rate at which such fixation occurs has thus been a major goal for evolutionary biology (McCandlish and Stoltzfus, 2014), as well as more applied biosciences, such as population management and conservation (Wright et al., 2009). The rate of genetic adaptation of a diploid population is often quantified with 2*N*_T_*µP*_fix_ where *N*_T_ is the population size, *µ* is the per-generation per-locus probability that a beneficial mutation occurs, and *P*_fix_ is the probability that it fixes (Kimura, 1962, 1968; Kimura and Ohta, 1971; Kryazhimskiy and Plotkin, 2008; Lanfear et al., 2014). This assumes that the mutation rate *µ* is small, such that the time taken for a beneficial allele to fix is negligible compared to the waiting time before such a mutation arises. Nevertheless, the time a beneficial allele takes to fix is in some cases relevant as it scales comparably to the number of generations it takes to arise, thus causing a slowdown in adaptation (e.g., in a large panmictic population, the fixation time of a mutation causing a fecundity advantage *s* ≪ 1 scales with log(2*N*_T_*s*)/*s*, which compared with the expected time the mutation takes to arise, 1/[2*N*_T_*µP*_fix_] with *P*_fix_ = 2*s*, entails that fixation time is increasingly relevant as 2*N*_T_*µ* log(2*N*_T_*s*) increases; Weissman and Barton, 2012; Charlesworth, 2020, 2022; for more general considerations, see Hendry and Kinnison, 1999). Additionally, because whether an allele fixes quickly or slowly influences the genetic signatures of adaptation at surrounding loci, the time to fixation may be important in the detection of selected sites in the genomes of natural and experimental populations (Messer and Petrov, 2013; Charlesworth, 2020, 2022).

The probability that a beneficial mutation fixes and the time it takes to do so both depend on an interplay between selection and genetic drift. This interplay is especially well understood under panmixia, i.e. where individuals mate and compete at random (Crow and Kimura, 1970; Ewens, 2004). In particular, because a rare allele is found almost exclusively in heterozygotes under panmixia, the probability of fixation of a single-copy *de novo* mutation strongly depends on its genetic dominance (or penetrance). Dominant beneficial alleles are more likely to fix than recessive ones as their effects are more immediately exposed to positive selection (Haldane, 1927a). Population size, which scales inversely with genetic drift, increases fixation time (Kimura and Ohta, 1969), but tends to have limited effects on the probability that a newly arisen mutation will fix (Kimura, 1962). In fact, the probability of fixation of a *de novo* mutation becomes independent from population size in the limit of infinite population size such that invasion can be modelled as a branching process and invasion implies fixation (Haldane, 1927b; Otto and Whitlock, 2001).

Genetic adaptation is not restricted to the fixation of *de novo* mutations, but can also stem from standing genetic variation (Orr and Betancourt, 2001; Hermisson and Pennings, 2005; Pennings and Hermisson, 2006a,b; Barrett and Schluter, 2008; Hermisson and Pennings, 2017). This standing variation is thought to be neutral or mildly deleterious until an environmental change takes place such that it becomes beneficial. The probability that such a variant fixes and the time it takes to do so are especially sensitive to its initial frequency, with greater frequency increasing the probability of fixation (Kimura, 1962) and reducing the time taken to fix (Kimura and Ohta, 1969). The initial frequency, in turn, depends on how selection and genetic drift shaped variation before it turned beneficial, which has also been extensively studied in well-mixed populations (Kimura et al., 1963; Crow and Kimura, 1970; Orr and Betancourt, 2001; Ewens, 2004).

Many natural populations, however, are not well-mixed. The physical constraints on movement often cause dispersal to be limited, leading to genetic structure through limited gene flow (Clobert et al., 2001). Such genetic structure is extremely widespread though often mild with many estimates of among-populations genetic differentiation *F*_ST_ of the order of 0.1 (e.g. Ståhl, 1981; Glover et al., 2013; Benzie, 2000; Giles and Goudet, 1997; Potenko and Velikov, 1998; Tamaki et al., 2008; Irvin et al., 1998; Kumar and Singh, 2017; Forstmeier et al., 2007; pp. 302-303 in Hartl and Clark, 2007 for an overview). Genetic structure influences both drift and selection as it modulates effective population size *N*_e_ (Wang and Caballero, 1999; Rousset, 2004), and generates kin selection and inbreeding (Hamilton, 1964; Frank, 1998; Rousset, 2004; Charlesworth and Charlesworth, 2010). The influence of inbreeding on genetic adaptation can be investigated independently by considering populations where selfing (or assortative mating) takes place but that are otherwise well-mixed (so that there is no kin selection or competition, Glémin and Ronfort, 2013; Newberry et al., 2016; Hartfield and Bataillon, 2020; Charlesworth, 2020). These investigations show that selfing tends to speed up fixation as it causes both: (i) an increase in homozygosity that exposes rare recessive alleles more readily to selection; as well as (ii) a decrease in effective population size that reduces the time to fixation.

How limited dispersal affects the probability of fixation through selection and drift is well-studied in the island model of dispersal, showing for instance that the fixation probability of beneficial alleles is increased by limited dispersal when recessive and decreased when dominant (Roze and Rousset, 2003; Whitlock, 2003; Rousset, 2004). Assuming that dispersal between demes is so rare that segregation time within demes can be ignored, Slatkin (1981) shows that limited dispersal always increases the time to fixation of *de novo* mutations. Using the diffusion approximation and thus considering segregation time within demes, Whitlock (2003) also reports that limited dispersal makes the total time to fixation increase, through an increase in *N*_e_ as well as in kin competition (p. 778 in Whitlock, 2003). Meanwhile, the implications of limited dispersal for the time taken by standing genetic variants to fix remain understudied (though see Paulose et al., 2019 for a discussion on this under isolation by distance).

Here, we contribute to this literature by investigating the rate of fixation of *de novo* and standing variants in subdivided populations. Firstly, we revisit the time to fixation of *de novo* mutations, complementing the analysis found in Whitlock (2003). We show that limited dispersal can in fact speed up fixation of non-additive alleles, as long as selection is not too weak and dispersal is mildly limited such that it generates *F*_ST_ < 0.2, which is typical of many natural populations. Secondly, we combine the waiting time and time to fixation to investigate the total rate of fixation from *de novo* mutations (as done in Glémin and Ronfort, 2013 for selfing). Thirdly, we investigate the impact of limited dispersal on the time for standing variation to fix. Finally, we consider the influence of meta-population dynamics whereby subpopulations can go extinct and be recolonized.

## 2 Model

### 2.1 Life cycle, genotype and fecundity

We consider a monoecious diploid population that is subdivided among *N*_d_ demes, each carrying a fixed number *N* of adult individuals (so that the total population size is *N*_T_ *= N*_d_ · *N*). Generations are discrete and non-overlapping with the following life cycle occurring at each generation: (i) Each adult produces a large number of gametes according to its fecundity and then dies. (ii) Each gamete either remains in its natal deme with probability 1 − *m* or disperses with probability *m*. We assume that *m* > 0 so that demes are not completely isolated from one another. Dispersal is uniform among demes, following the island model (Wright, 1931). (iii) Finally, gametes fuse randomly within each deme to form zygotes that in turn compete uniformly within each deme for *N* breeding spots to become the adults of the next generation.

We are interested in evolution at an autosomal locus where two alleles segregate: a wild-type allele *a* and a beneficial mutant allele *A*. An individual’s genotype determines its fecundity. As a baseline, *aa* individuals have a fecundity of 1, while relative to this, *Aa* and *AA* individuals have fecundity of 1 +*hs* and 1 + *s*, respectively. The parameters 0 ≤ *h* ≤ 1 and *s* > 0 thus capture the dominance and selective effects of *A*, respectively. For simplicity, we assume throughout the main text that selection is soft, i.e. that the same total number of gametes is produced in each deme. The case of hard selection is explored in our Supplementary Material (File S1) where we show that our main results are not affected by whether selection is soft or hard.

### 2.2 Diffusion approximation

The dynamics of the frequency *p* of the allele *A* in the whole population can be approximated by a diffusion process under the island model of dispersal (e.g. Barton, 1993; Roze and Rousset, 2003; Whitlock, 2003; Cherry and Wakeley, 2003; Cherry, 2003; Wakeley, 2003; Wakeley and Takahashi, 2004; Lessard, 2009; note that one cannot follow a single allele frequency when the population experiences isolation by distance, see Rousset, 2004 for general considerations). We follow the framework developed in Roze and Rousset (2003), which assumes that selection is weak and that the number *N*_d_ of demes is large (i.e. *s* ∼ 𝒪(*δ*) and *N*_d_ ∼ 𝒪(1/*δ*) where *δ* > 0 is small). If, in addition, demes are large and dispersal is weak (i.e. *N* → ∞ while *m* → 0 such that the number of immigrants *Nm* is of order 1), then allelic segregation within demes also follows a diffusion process (e.g. Whitlock, 2003). Here, we will in general allow for *m* to be arbitrarily large to investigate deviations from panmixia (i.e. from *m =* 1). The diffusion approximation is based on the expectation and variance in the change in *p*, which we describe below.

#### 2.2.1 Expected frequency change

We show in section A.1 in File S1 that the expected change in allele frequency *p* can be written as,

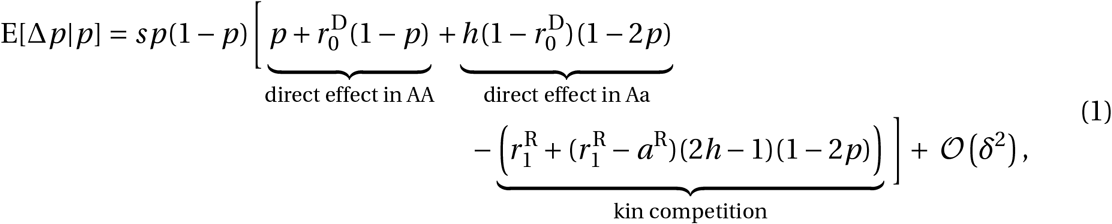

where 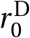 is the probability that the two homologous genes of an individual are identical-by-descent (IBD); 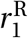 is the probability that two genes sampled from the same deme with replacement are IBD; *a*^R^ is the probability that two homologous genes of an individual, plus a third gene sampled from the same deme at random, are all IBD. These three coalescent probabilities are computed under neutrality (i.e. with *δ =* 0) and their expression in terms of dispersal and population size can be found in Table 1 (see section A.3 in File S1 for derivations). Equation (1) is equivalent to eq. (23) of Roze and Rousset (2003) after plugging in their fitness eq. (36) (and additionally using our eq. A24 from File S1).

**Table 1:** Probabilities of coalescence in the island model. Expressions for the various probabilities of coalescence that are relevant to the analysis, including their values in the limit of low dispersal and large patches (*m* → 0 and *N* → ∞ such that *Nm* remains constant, see section A.3 in File S1 for derivations). 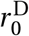 is the probability that the two homologous genes of the same individual are identical-by-descent (IBD), which is equivalent to *F*_IT_ in the island model of dispersal; 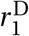 is the probability that two different genes sampled from the same deme are IBD, which is equivalent to *F*_ST_ in the island model of dispersal; 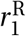 is the probability that two genes sampled from the same deme with replacement are IBD; and *a*^R^ is the probability that two homologous genes of an individual coalesce with a third gene sampled from the same deme at random (Roze and Rousset, 2003; Rousset, 2004).

| Symbol | Expression | Limit |
| --- | --- | --- |
| $r_0^D$ | $\frac{(1-m)^2}{2N-(1-m)^2(2N-1)}$ | $\frac{1}{1+4Nm}$ |
| $r_1^D$ | " | " |
| $r_1^R$ | $\frac{1}{2N} + \left(1 - \frac{1}{2N}\right) r_1^D$ | " |
| $a^R$ | $\frac{1}{N} r_0^D + \left(1 - \frac{1}{N}\right) \frac{\left[1 + 3(2N-1)r_1^D\right](1-m)^3}{(2N)^2 - (2N-1)(2N-2)(1-m)^3}$ | $\frac{1}{(1+4Nm)(1+2Nm)}$ |

Equation (1) decomposes selection on allele *A* among three effects. These can be understood by first considering that in a well-mixed population (so that 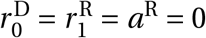), eq. (1) reduces to: E[Δ*p*|*p*] *= sp*(1 − *p*) *p* +*h*(1 − 2*p*) + 𝒪(*δ*^2^) (Crow and Kimura, 1970, p. 408). In this baseline expression, the first term within square brackets, *p*, captures selection on *A* owing to the effects of the allele on the fitness of its bearer in homozygotes, while *h*(1 − 2*p*) captures selection through heterozygotes. When dispersal is limited, direct effects increase to 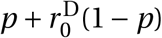 through homozygotes and decrease to 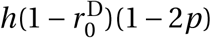 through heterozygotes in eq. (1). Selection through homozygotes is therefore more important under limited dispersal. This is because mating within demes leads to inbreeding and therefore a relative excess of homozygotes and a deficit of heterozygotes (according to 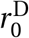).

The remaining terms of eq. (1) – labelled “kin competition” – capture a second effect of limited dispersal: that competing individuals are more likely to carry identical gene copies than randomly sampled individuals. As shown by the negative sign in front of these terms in eq. (1), kin competition decreases selection on beneficial alleles. This is because kin competition results in an individual’s reproductive success coming at the expense of genetic relatives. Equation (1) further shows that for additive alleles (*h =* 1/2), kin competition effects scale with genetic differentiation 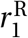 only. For non-additive alleles, however, these effects also depend on allele frequency *p*, with kin competition effects being stronger when dominant alleles are rare (*h* > 1/2 and *p* < 1/2) or when recessive alleles are common (*h* < 1/2 and *p* > 1/2).

In the limit of low dispersal and large demes (*m* → 0 and *N* → ∞), the pairwise probabilities of coalescence 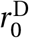 and 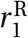 are equal to *F*_ST_ (Rousset, 2004) where *F*_ST_ = 1 (1 + 4*Nm*) (Wright, 1931), while the threeway probability of coalescence can be written as 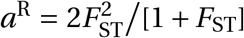 (Whitlock, 2002). As a result, eq. (1) can be expressed as

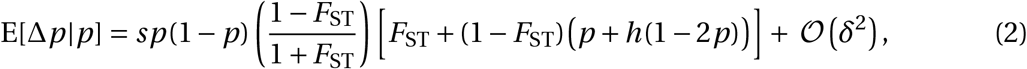

which is the same as eq. (12) of Whitlock (2002) (with his *η =* 0; we compare the times to fixation computed with eq. 2 and with eq. 1 in Figure A in File S1, which shows overall good agreement between the two with *N =* 100).

#### 2.2.2 Variance in frequency change and effective population size

The variance in allele frequency change can be written in the form

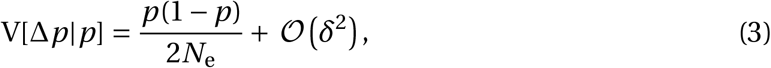

where effective population size *N*_e_ for our model is given by eq. (28) of Roze and Rousset (2003),

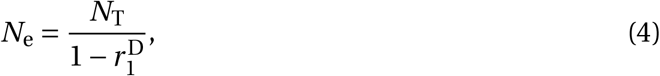

with 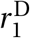 is the probability that two different genes sampled from the same deme are IBD. See section A.2 in File S1 for derivation of eq. (4) and Table 1 for the probability 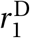 in terms of deme size and dispersal rate. The effective population size of eq. (4) can also be written in the low dispersal and large demes limit, such that 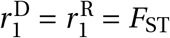. This substitution leads to the classical expression, *N*_e_ *= N*_T_/(1 −*F*_ST_) or *N*_e_ *= N*_T_ [1 + 1/(4*Nm*)] (Wright et al., 1939; Whitlock and Barton, 1997; Whitlock, 2003; Roze and Rousset, 2003).

It may be useful to consider the scaled or “effective” selection gradient in the low dispersal and large demes limit, which reads as

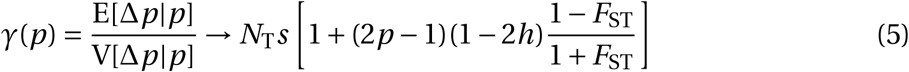

(using eq. 2 - 4). Two points are worth mentioning from eq. (5). First, it shows that it is relevant to scale the fecundity advantage *s* with the total population size *N*_T_ when comparing the strength of selection among treatments. Second, it makes clear how effective selection depends on an interaction between allele frequency *p* and genetic dominance *h* that is modulated by gene flow *F*_ST_. In a well-mixed population such that *F*_ST_ = 0, *γ*(*p*) is greater when dominant alleles are rare (*h* > 1/2 and *p* < 1/2) or when recessive alleles are common (*h* < 1/2 and *p* > 1/2) compared to *γ*(*p*) for an additive allele (*h =* 1/2). By creating to an excess of homozygosity, limited gene flow mitigates these differences by a factor 0 ≤ (1 −*F*_ST_) (1 +*F*_ST_) ≤ 1.

### 2.3 Probability and time of fixation

From the scaled selection gradient *γ*(*p*), the probability of fixation *P*_fix_(*p*_0_) of *A* with initial frequency *p*_0_ is given by

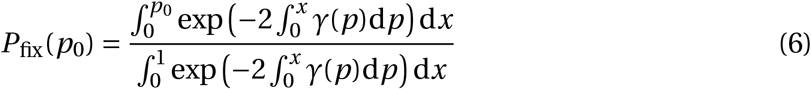

(Crow and Kimura, 1970, p. 424). The expected number *T*_fix_(*p*_0_) of generations that an allele takes to fix (conditional on its fixation) given that its frequency is *p*_0_ at generation *t =* 0 is,

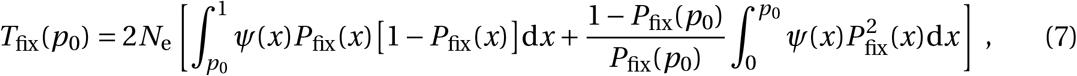

with

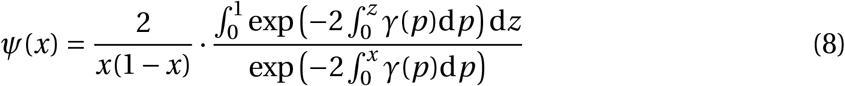

(Crow and Kimura, 1970, p. 430 with their *ψ*(*x*) scaled by 1/(2*N*_e_)).

Equation (7) highlights how the time to fixation scales with 2*N*_e_ (the “coalescent timescale”; Kimura, 1962; Charlesworth, 2020), which depends on limited dispersal (eq. 4). There are therefore two main pathways for limited dispersal to influence the time to fixation: (i) by modulating the scaled selection gradient for non-additive alleles (as seen most clearly in eq. 5); and (ii) by boosting effective population size (eq. 4). To investigate this interaction, eqs. (6)-(8) were numerically integrated with R using the built-in function `integrate` (with maximum number of subdivisions increased to 100^*′*^000). The time to fixation for non-additive alleles (*h≠*1/2) involves an integral with an integrand that spans many orders of magnitude, which can be prone to instability during numerical analysis. In the case of report of bad integrand behaviour, we translated integration limits by a small amount *ϵ* that we kept as low as possible (*ϵ* ≤ 10^−5^ always).

To calculate the time to fixation more straightforwardly, Charlesworth (2020) suggests an approximation based on a decomposition of fixation dynamics between three phases: two stochastic phases when *p* < *p*_1_ and *p* > *p*_2_ connected by a deterministic phase for *p*_1_ ≤ *p* ≤ *p*_2_ (following semi-deterministic approaches, e.g. Martin and Lambert, 2015). This approximation can be summarized as

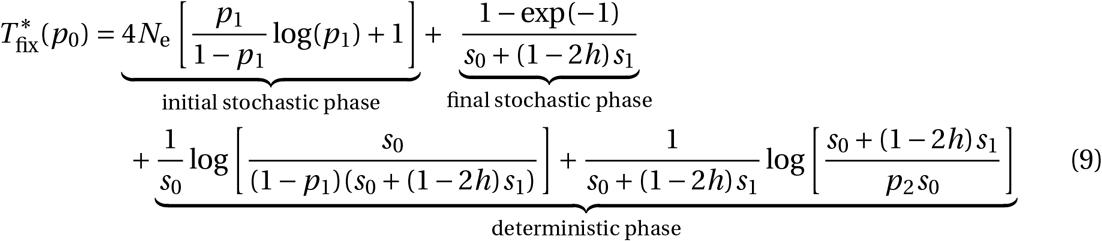

(“initial stochastic phase” is on p. 15, “final stochastic phase” in eq. A7, and “deterministic phase” in eq. 6b of Charlesworth, 2020), where

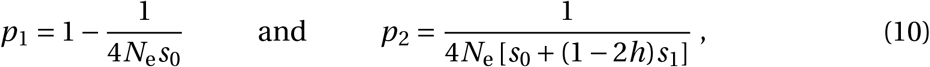

and *s*_0_ and *s*_1_ are defined from decomposing the expected frequency change as E[Δ*p*|*p*] *= p*(1 − *p*) *s*_0_ + (1 − 2*h*)*s*_1_*p*, which by comparison with our eq. (1) yields,

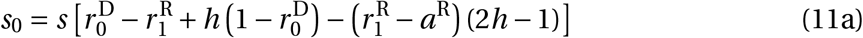

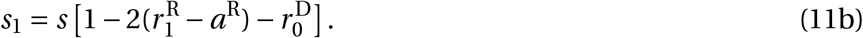

We will also use this approximation to compute fixation time.

## 3 Results

### 3.1 The antagonistic effects of limited dispersal on the time to fixation of *de novo* mutations

We first complement Whitlock (2003)’s analyses on the effects of limited dispersal on the time it takes for *A* to fix as a *de novo* mutation (i.e., when arising as a single copy, *P*_fix_(*p*_0_) with *p*_0_ = 1 (2*N*_T_)). Whitlock (2003) considers a fecundity advantage of *s =* 2 · 10^−4^ such that *N*_T_*s =* 2, and shows that in this case, limited dispersal always increases the time to fixation by increasing effective population size (due to the factor 2*N*_e_ in eq. 7; see also panel B in Figure A in File S1). We consider the case where selection is stronger, though still weak, where individuals that carry two copies of *A* experience a 1% increase in fecundity (*s =* 0.01) in a population of 200 demes of 100 individuals (such that with *N*_T_*s =* 200 as in Roze and Rousset, 2003; we consider other selection strengths later). We show that by modulating the interaction between selection and drift, limited dispersal can decrease the time to fixation in this case.

We integrated eq. (7) with *p*_0_ = 1/(2*N*_T_) for a range of dispersal *m* and dominance *h* values. Results of these calculations and of individual based simulations are shown in Figure 1. We find that the effect of limited dispersal on the time to fixation depends on the dominance of the beneficial allele *A*. Where dominance is incomplete (approximately 0.1 ≤ *h* ≤ 0.9, Figure 1B), the expected time to fixation always increases as dispersal becomes more limited (Figure 1A, blue). In contrast, the time to fixation of a partially dominant (*h* > 0.9) or partially recessive (*h* < 0.1) allele initially decreases as dispersal becomes limited and only increases once past below a dispersal threshold (Figure 1A, green and purple). The reason the time to fixation eventually increases when dispersal becomes severely limited (approx. *Nm* < 1 so that *F*_ST_ > 0.2 in Figure 1A) is because *N*_e_ increases hyperbolically as dispersal decreases (recall *N*_e_ ∼ *N*_T_ [1 + 1/(4*Nm*)], see below eq. 4). As a result, the effects of *N*_e_ on time to fixation overwhelms any other effects when dispersal becomes small. These results so far are consistent with those in Whitlock (2003), where selection is sufficiently weak such that the effects of limited dispersal on the time to fixation are mostly through its effects on *N*_e_ (recall eq. 7).

**Figure 1:**
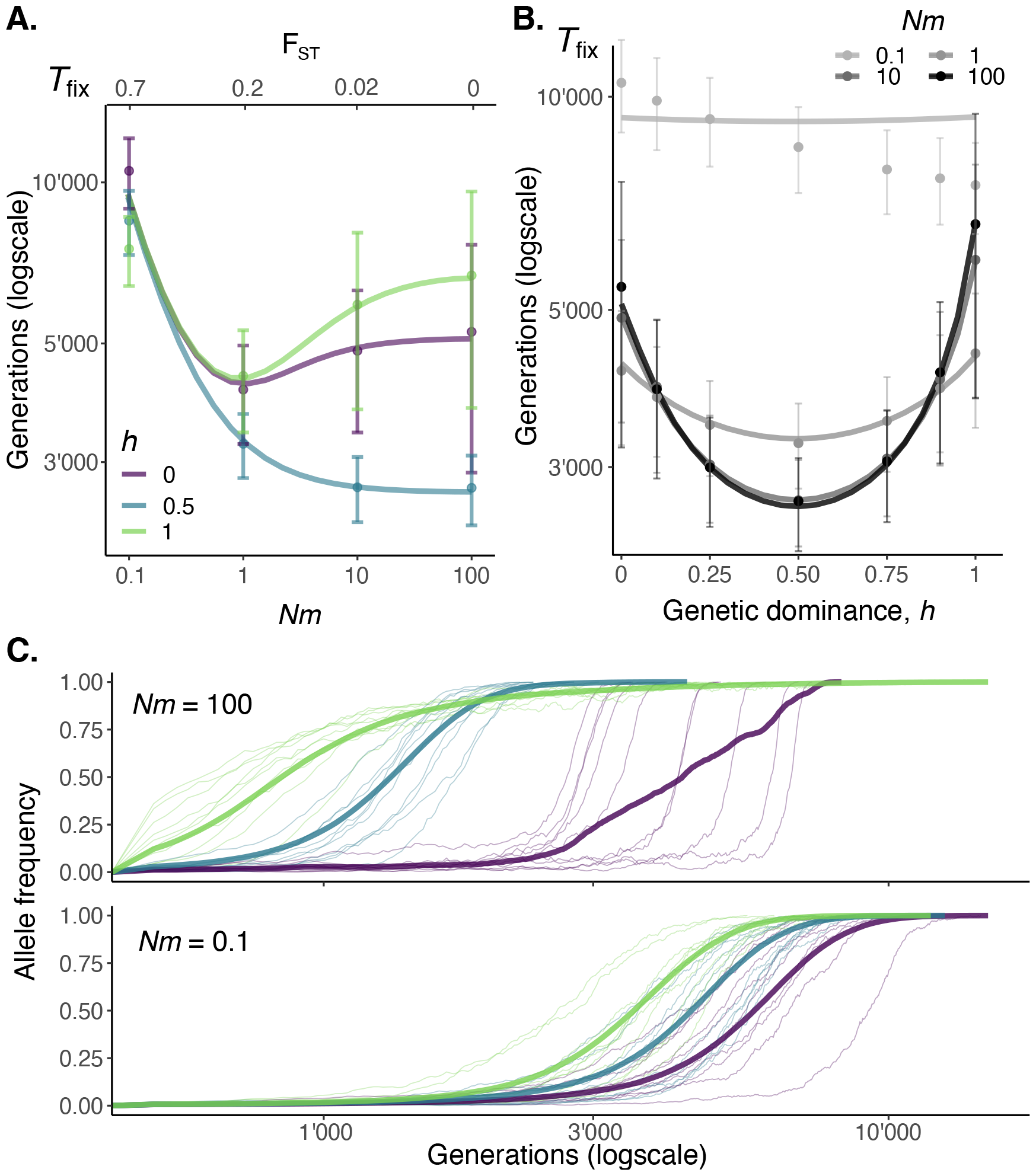
Time and trajectory to fixation of beneficial mutations according to dispersal. **A**. Expected time to fixation *T*_fix_(*p*_0_) of recessive (*h =* 0), additive (*h =* 0.5) and dominant (*h =* 1) alleles (in different colors, see legend) arising as single copies *p*_0_ = 1/(2*N*_T_), with solid lines from diffusion approximation (i.e. numerical integration of eq. 7) and dots as averages from simulations (fixation events among 40’000 replicates for each set of parameters, error bars show standard deviation, section B in File S1 for details). Parameters: *N*_d_ = 200, *N =* 100, *s =* 0.01. Under strong dispersal limitation, *Nm* ≲ 0.1, the time to fixation asymptotically approaches the neutral expectation *T*_fix_ ∼ 4*N*_e_ ∼ 4*N*_T_[1 + 1/(4*Nm*)] (Kimura and Ohta, 1969), regardless of genetic dominance. **B**. Expected time to fixation *T*_fix_(*p*_0_), i.e. same as A, but plotted against dominance for different levels of dispersal (see legend). **C**. Fixation trajectories of beneficial mutations in a well-mixed (top, *Nm =* 100) and dispersal-limited (bottom, *Nm =* 0.1) population. For each level of dominance (in different colours, see A for legend), thin lines show ten randomly sampled trajectories, thick lines show the mean trajectory among all trajectories. Parameters: same as A.

Under mild dispersal limitation (approx. 1 < *Nm* < 100 such that *F*_ST_ ≤ 0.2), however, our results show that the increase in effective population size *N*_e_ can be outweighed by an increase in selection, resulting in partially recessive and partially dominant alleles fixing more rapidly than under panmixia (Figure 1A, dark gray line in B). The reason selection reduces the time to fixation here is because limited dispersal leads to inbreeding and thus a relative excess of homozygotes. How this excess boosts selection depends on whether the allele is recessive or dominant, as revealed by considering an increase in 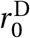 in eq. (1). For a recessive beneficial allele *A* (*h* < 1/2), selection on *A* is greater when *A* is relatively rare (i.e. *p* < 1/2) because in this case, inbreeding creates a relative excess of *AA* homozygotes through which the recessive allele *A* can be picked up by selection. For a dominant beneficial allele *A* (*h* > 1/2), the excess of homozygosity boosts selection at high frequency (i.e. *p* > 1/2) as it allows to purge more efficiently the deleterious (and recessive) *a* allele through an excess of *aa* individuals. These frequency-dependent effects are amplified by kin competition (the fact that under limited dispersal, competing individuals are more likely to carry identical gene copies than randomly sampled individuals, Rousset, 2004). Kin competition weakens selection as it results in an individual’s reproductive success coming at the expense of genetic relatives. The strength of kin competition on a non-additive allele depends on its frequency (see eq. 1). Specifically, kin competition is weaker and thus selection is stronger when a recessive allele is rare (*h* < 1/2 and *p* < 1/2) and a dominant allele is common (*h* > 1/2 and *p* > 1/2).

The frequency-dependent effects of limited dispersal on selection are reflected in the trajectory profiles of recessive and dominant alleles that fix (Figure 1C). In a panmictic population, a recessive beneficial allele tends to spend longer periods at low frequency (for enough homozygotes to appear) and a dominant allele at high frequency (for heterozygotes to be purged, Figure 1C, top). Under limited dispersal, however, these differences are mitigated as selection is increased at low frequency for recessive alleles and at high frequency for dominant alleles (e.g. eq. 5). As a result, the trajectory profiles of beneficial alleles that eventually fix become independent of their dominance as dispersal becomes limited (Figure 1C, bottom). This can also be seen from the decomposition of the time to fixation into three relevant phases according to the semi-deterministic approximation eq. (9). As shown in Figure 2, limited dispersal decreases the share of the time that recessive alleles spend in the initial stochastic phase (lower shaded region in top row of Figure 2) and the share that dominant alleles spend in the final stochastic phase (top shaded region in bottom row of Figure 2). In fact, owing to an excess homozygosity that reduces boundary effects at low and high frequency, the semi-deterministic approximation eq. (9) performs better under limited dispersal when alleles are recessive or dominant (compare black full and dashed gray lines in Figure 2).

**Figure 2:**
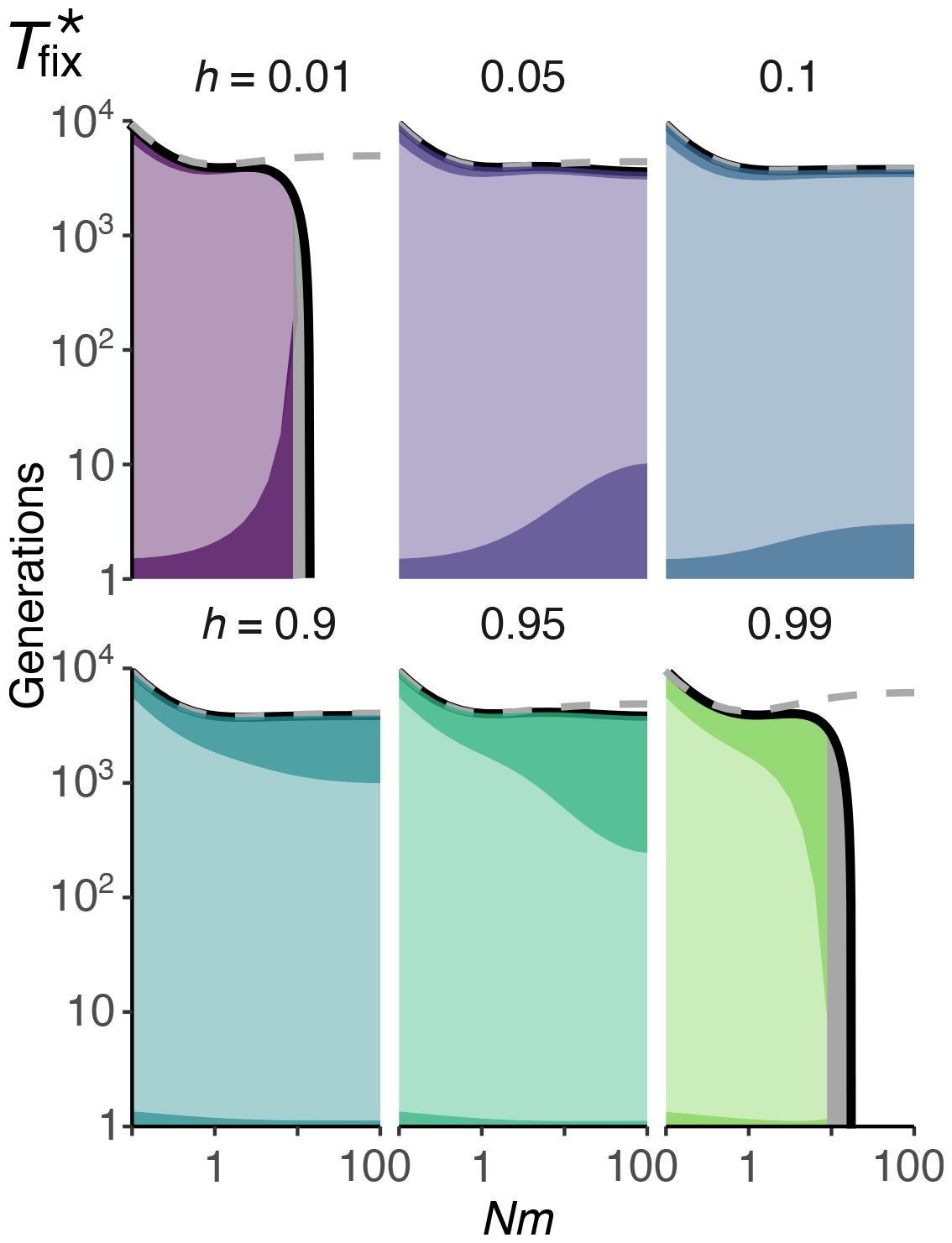
Semi-deterministic approximation to fixation time under limited dispersal. Solid black lines show Charlesworth (2020)’s approximation to the expected time to fixation 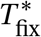 (eq. 9) of partially recessive (*h =* 0.01, 0.05, 0.1; top row) and partially dominant (*h =* 0.9, 0.95, 0.99; bottom row) alleles arising as single copies *p*_0_ = 1 /(2*N*_T_); and dashed gray lines show numerical integration of eq. (7). Shaded regions below curve represent the proportion of time spend in each phase of the approximation of eq. (9), from bottom to top: initial stochastic phase (dark shade), deterministic phase (light shade), final stochastic phase (dark shade). The shaded gray areas in top left (*h =* 0.01) and bottom right (*h =* 0.99) graphs indicate where eq. (9) diverges. Other parameters: same as Figure 1.

The above shows that limited dispersal can reduce the time to fixation of partially recessive and dominant alleles, provided the selection coefficient is above some threshold. We investigate this threshold numerically in panel C in Figure A in File S1, which suggests that it is close to *N*_T_*s =* 50 (e.g. such that carrying two copies of the beneficial allele causes a 0.25% increase in fecundity with *N*_d_ = 200 and *N =* 100). This value sits well within empirically estimated distribution of fitness effects (Eyre-Walker and Keightley, 2007).

Although the time to fixation is useful for multiple reasons (e.g. Whitlock, 2003; Glémin and Ronfort, 2013; Charlesworth, 2020, 2022), it may not always provide a good reflection of the time for a population to show high mean fecundity, especially when beneficial alleles are dominant. To see this, we computed the expected time taken for the genetic load to drop to 10% in the whole population (*τ*_G_), as well as within each deme (*τ*_L_), using individual based simulations (Figure 3). We observe *τ*_G_ < *τ*_L_ < *T*_fix_ throughout, with their differences larger the greater *h* is. This is because dominant alleles confer high fitness in heterozygotic form and thus allow the population to show low load at lower frequency. Therefore, though limited dispersal reduces the time to fixation of both recessive and dominant beneficial mutations, the time taken for the population to show high fecundity is reduced only for recessive alleles.

**Figure 3:**
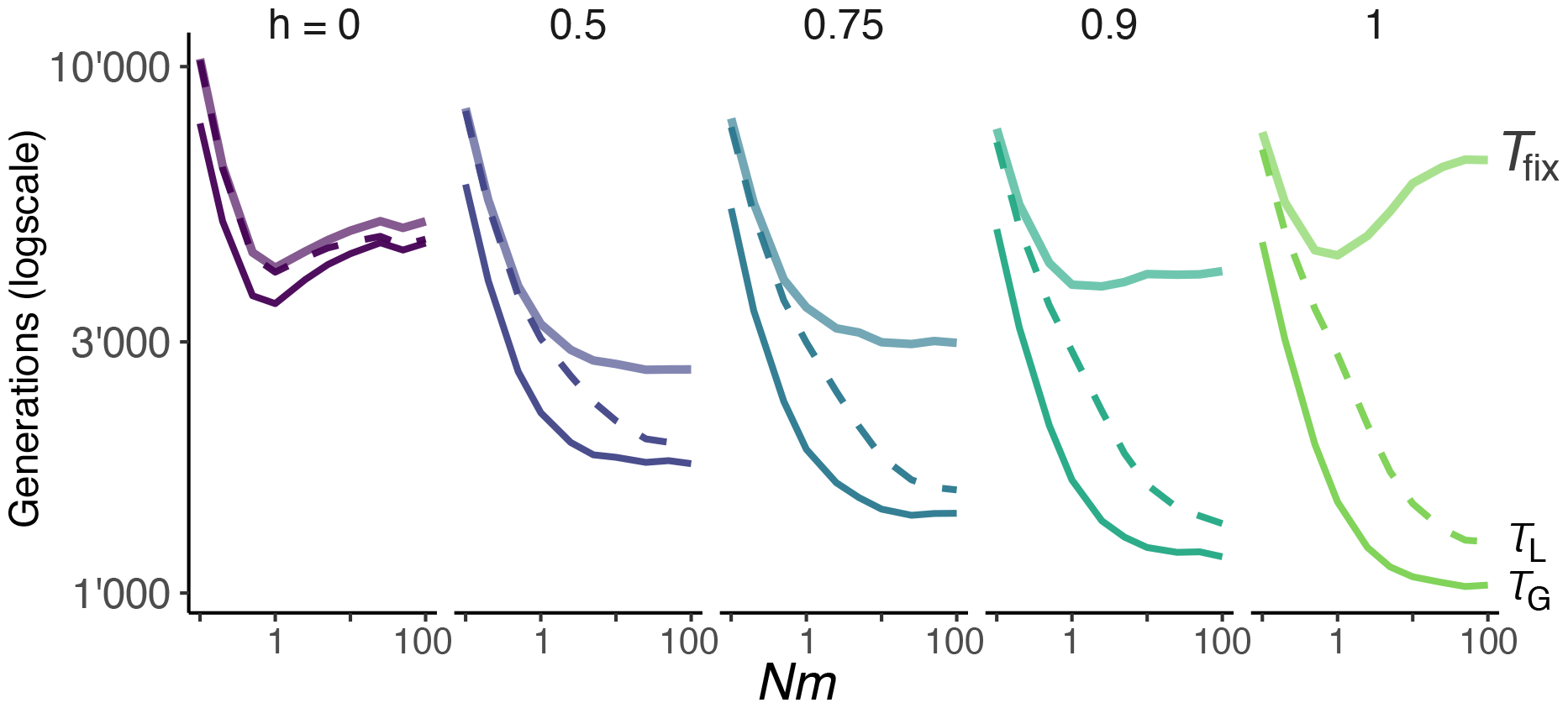
Expected time to purge 90% of the genetic load. We define the genetic load as *L =* (1 + *s* − *z*) / *s*, i.e. as the difference between the population mean fecundity, *z* (eq. A10 in File S1), and the maximum fecundity, 1 + *s*, normalized such that a population monomorphic for the wild-type allele, *aa* (*z =* 1), has *L =* 1, while a population where the beneficial mutation *A* has fixed has no genetic load, *L =* 0. We also define a local genetic load *L*_*i*_ in each deme *i* as *L*_*i*_ = (1 + *s* − *z*_*i*_)/*s*, where *z*_*i*_ is the mean fecundity at deme *i* (eq. A8). Plots show the average time to fixation *T*_fix_ (thick line), average time to purge 90% of the load *τ*_G_, i.e. average time for *L =* 0.1 (thin line), and average time to purge 90% of genetic load in every deme *τ*_L_, i.e. average time for max_*i*_ *L*_*i*_ = 0.1 (dashed line), for recessive (*h =* 0), additive (*h =* 0.5) and (partially) dominant (*h =* 0.75, 0.9 and 1) alleles (in different columns) arising as single copies *p*_0_ = 1/(2*N*_T_) in individual-based simulations (section B in File S1 for details). Other parameters: same as Figure 1.

### 3.2 The impact of limited dispersal on the total time for *de novo* mutations to arise and fix

In addition to the number of generations taken for a beneficial allele to fix, another relevant consideration is how long it takes for such an allele to emerge. To capture this, we follow Glémin and Ronfort (2013) and quantify the total expected number *T*_new_ of generations for an adaptive *de novo* mutation to fix in the population by the sum of two terms:

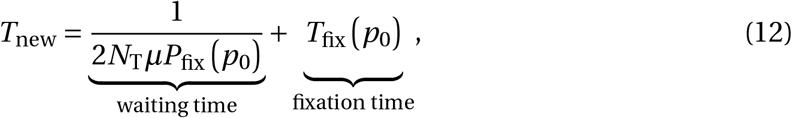

where *µ* is the mutation rate from *a* to *A*, and *P*_fix_ is the fixation probability of *A* when it arises as a single copy, i.e. when *p*_0_ = 1 (2*N*_T_) (eq. 6 for the diffusion approximation to this probability). The first term is the expected number of generations for the emergence of a beneficial mutation that fixes, and the second is the expected number of generations taken by such fixation. The underlying assumption behind using eq. (12) is that beneficial mutations appear at a per-site per-generation rate *µ* that is such that new mutations segregate independently (as in e.g. Gillespie, 1994’s strong-selection weak-mutation regime).

The waiting time is inversely proportional to the fixation probability *P*_fix_, whose dependence on limited dispersal is well known: while limited dispersal has no influence on the probability of fixation of additive alleles, it increases (respectively, decreases) the probability that a recessive (dominant) beneficial allele fixes (Roze and Rousset, 2003; Whitlock, 2003). Hence, the waiting time for a fixing additive allele is not affected by limited dispersal, but is reduced for a recessive allele and increased for a dominant allele (Figure 4A). Accordingly, the total number of generations *T*_new_ for fixation always increases with limited dispersal when such an allele is additive (as *T*_fix_ increases, section 3.1, Figure 4A central column). Recessive alleles, meanwhile, benefit from limited dispersal in two ways, as limited dispersal not only reduces the time to fixation (provided that dispersal is not too limited) but also the waiting time for such a fixing allele to appear. This results in a significant drop in the total time for fixation as dispersal becomes limited, before eventually increasing under severely limited dispersal (Figure 4A left column).

**Figure 4:**
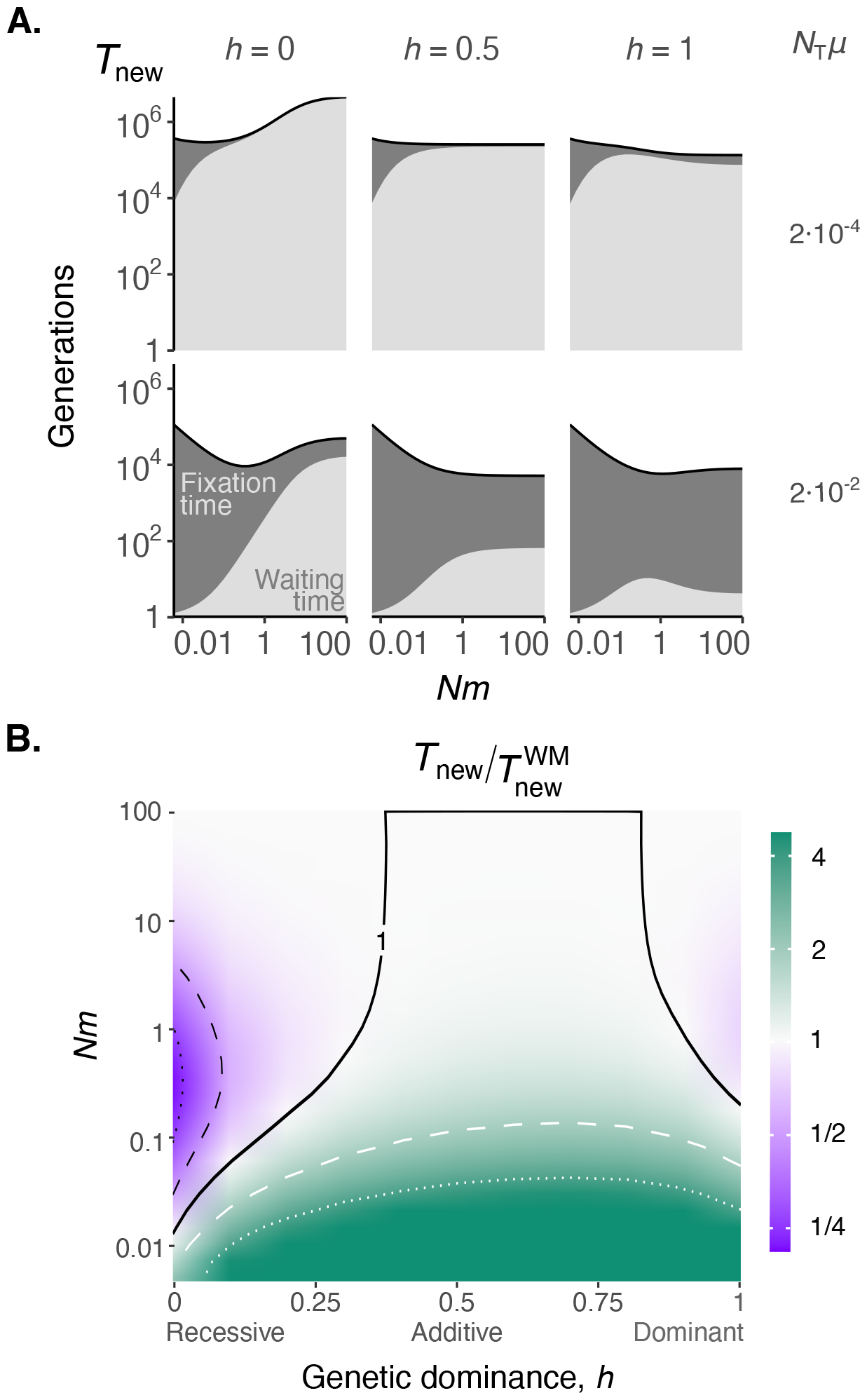
Total time before a *de novo* mutation arises and fixes under limited dispersal. **A**. Expected total time *T*_new_ of fixation (on a log scale) of a recessive (*h =* 0, left), additive (*h =* 0.5, middle) and dominant (*h =* 1, right) *de novo* mutation *p*_0_ = 1/ (2*N*_T_) for different mutation rates (*N*_T_*µ =* 2 · 10^−4^, top; *N*_T_*µ =* 2 · 10^−2^, bottom), with solid black lines from eq. (12) (with eqs. 6-7). Dark and light gray shades underneath curves represent the proportion of time spent in each component of *T*_new_ (on a linear scale). Parameters: same as Figure 1. **B**. Effect of population subdivision on the total time to fixation according to scaled dispersal rate *Nm* and genetic dominance *h*. Ratio between the expected time to fixation of *de novo* mutations under limited dispersal *T*_new_ and panmixia (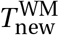, i.e. where *m =* 1, both from eq. 12 with eqs. 6-7). Full contour shows 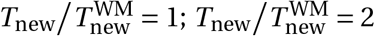 and 4 in black dashed and dotted, respectively; 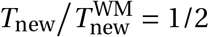 and 1/4 in white dashed and dotted, respectively. Parameters: *N*_T_*µ =* 2 · 10^−2^, other parameters: same as Figure 1.

The case of dominant alleles is more complicated as on one hand limited dispersal increases the waiting time, but on the other reduces the time to fixation (section 3.1 and Figure 4A right column). The balance between these opposing effects depends on the mutation rate *µ*. When this rate is very small, the waiting time dominates eq. (12) so that limited dispersal always increases the total time *T*_new_ for dominant alleles to fix (Figure 4A top-right). As the mutation rate increases, however, the time to fixation becomes more relevant in eq. (12) so that limited dispersal may reduce total time *T*_new_, though less than for recessive alleles (Figure 4A bottom-right).

Overall, we thus find that the rate of fixation of adaptive alleles depends on the interaction between the dominance *h* of these alleles and dispersal *m*. To see this more definitively, we compare *T*_new_ across levels of dispersal and dominance with *T*_new_ under panmixia in Figure 4B for a mutation rate of *N*_T_*µ =* 0.02. This figure shows that, for this set of parameters, the total time for the fixation of *de novo* mutations can be up to four times more rapid under limited dispersal compared to panmixia when beneficial alleles are recessive (Figure 4B, dark purple region, dotted black lines). The effect for dominant alleles, although weaker, is still non-negligible with fixation up to 30% faster under limited dispersal (Figure 4B, light purple region where *h* > 0.5). Below a dispersal threshold, however, fixation is slower whatever the dominance of beneficial alleles (Figure 4B, green region).

### 3.3 Fixation from standing variation: limited dispersal and dominance reversal

To investigate fixation from standing genetic variation, we now let the initial frequency of allele *A* in the whole global population *p*_0_ be a random variable, whose distribution is determined by assuming that *A* is initially deleterious, maintained at a mutation-selection-drift equilibrium until an environmental change takes place that causes *A* to become beneficial (following Orr and Betancourt, 2001; Hermisson and Pennings, 2005, 2017; Orr and Unckless, 2008; Glémin and Ronfort, 2013). Given a realisation *p*_0_, the initial frequency in each deme when environment changes is thus on average *p*_0_ but there is variation among demes, i.e. there is genetic differentiation among demes due to local sampling effects. The expected number of generations taken for fixation is now computed as

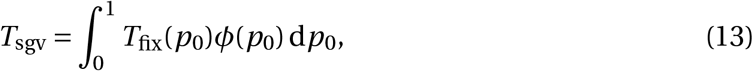

where *φ*(*p*_0_) is the probability density function for the frequency *p*_0_ of allele *A* in the whole population at the time of the environmental change when allele *A* becomes beneficial. This distribution *φ*(*p*_0_) can be calculated at mutation-selection-drift equilibrium using the diffusion approximation (eq. A49 in Supplementary Material). Prior to the environmental change, allele *A* is deleterious such that *aa, Aa* and *AA* individuals have fecundity of 1, 1−*h*_D_*s*_D_, and 1−*s*_D_, respectively, while mutations from *a* to *A*, and from *A* to *a*, occur at rate *µ*. Population structure and dispersal rate are assumed to be the same before and after the environmental change. We computed eq. (13) numerically under different values of genetic dominance before (*h*_D_) and after (*h*) the environmental change and various dispersal rates (*m*).

Let us first consider scenarios where the dominance of *A* is preserved before and after the environmental change (i.e. *h*_D_ *= h*). We find that under mildly limited dispersal, fixation takes longer when *A* is additive and shorter time when *A* is dominant (Figure 5A blue and green). Thus, limited dispersal has the same effect on the time for *A* to fix as a standing genetic variant than on the total time *T*_new_ for *A* to fix as a *de novo* mutation (provided mutation is strong enough so that waiting time does not dominate *T*_new_ when *A* is dominant). By contrast, whereas limited dispersal can lead to shorter time *T*_new_ for *A* to fix as a *de novo* mutation when recessive, it always increases the time *T*_sgv_ for *A* to fix as a standing genetic variant (Figure 5A purple). To understand this, recall that when the frequency of a beneficial recessive allele *A* is low, dispersal limitation speeds up the segregation of that allele by producing an excess of homozygotes *AA*. If such an allele is initially deleterious and recessive, however, its initial frequency *p*_0_ tends to be higher at the moment of environmental change. Consequently, the allele is likely to already exist in the homozygous form when it becomes beneficial, and thus is easily picked up by selection regardless of limited dispersal (Figure 5B purple for the distribution *φ*(*p*_0_)). Similarly to *de novo* mutations, the sweeping trajectories of standing genetic variants with different levels of dominance also become more similar as dispersal is reduced, in fact converging to the trajectories of recessive alleles under panmixia (compare top and bottom of Figure 5C).

**Figure 5:**
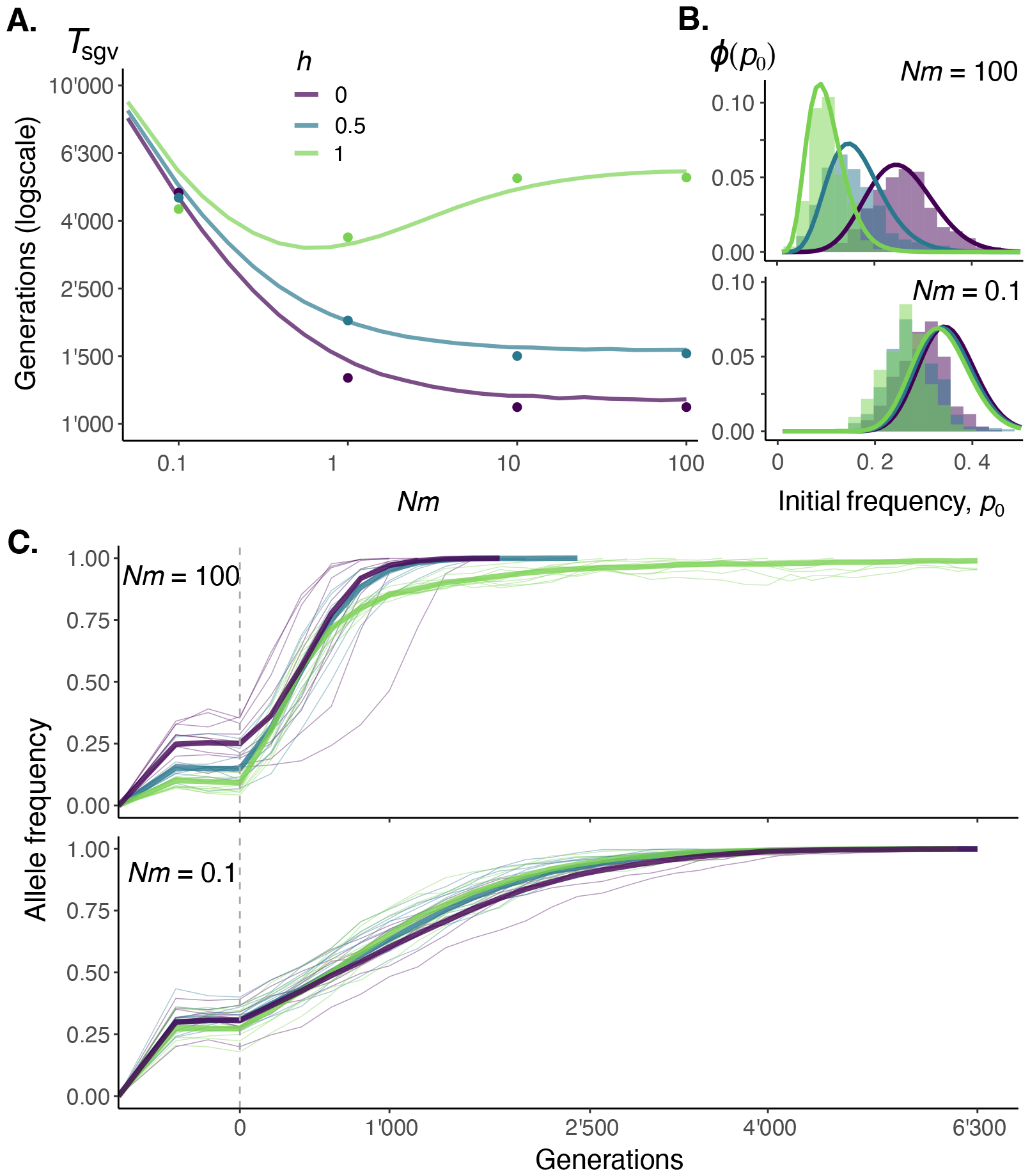
Fixation times of standing genetic variants. **A**. Expected time *T*_sgv_ of fixation (on a log scale) of a standing recessive (*h =* 0), additive (*h =* 0.5) and dominant (*h =* 1) variants (in different colors, see legend) with solid lines from diffusion approximation (eq. 13 with eqs. 7 and A49), and dots for the average from individual based simulations (300 replicates for each set of parameters, section B in File S1 for details; we do not show standard deviations of simulations here as they are typically large and therefore lead to an overcrowded figure; this is because *T*_sgv_ is affected by two sources of variance: variance in *p*_0_ and in the time to fixation). Parameters: *s*_D_ = 10^−3^, *N*_T_*µ =* 2, *h*_D_ *= h*, other parameters: same as Figure 1. **B**. Distribution of initial frequencies *φ*(*p*_0_) at the moment of environmental change in a well-mixed (top, *Nm =* 100) and dispersal-limited (bottom, *Nm =* 0.1) population. Vertical bars represent histograms of simulations and lines from diffusion approximation (eq. A49 in File S1). Note that the diffusion approximation fares less well when *Nm =* 0.1 as dispersal is much weaker than selection (Roze and Rousset, 2003; Wakeley, 2003). Parameters: same as A. **C**. Fixation of standing genetic variants in a well-mixed (top, *Nm =* 100) and a dispersal-limited (bottom, *Nm =* 0.1) population. Environmental change takes place at *t =* 0 (dashed vertical line). For each level of dominance (in different colours, see A for legend), thin lines show ten randomly sampled trajectories, thick lines show the mean trajectory among all trajectories. Parameters: same as A.

One scenario that has been argued to be particularly relevant in the context of fixation from standing genetic variation is that of dominance reversal, whereby an initially recessive deleterious allele (*h*_D_ = 0) becomes beneficial and dominant (*h =* 1) in the new environment (Muralidhar and Veller, 2022). This facilitates fixation because at mutation-selection-drift equilibrium, a recessive deleterious allele can be maintained at significant frequency, such that it can be readily picked up by selection when it turns beneficial, especially if it simultaneously becomes dominant. Comparing the case where *A* is additive before and after the environmental change, with the case where it shifts from being recessive to dominant, we see that limited dispersal reduces the effects of dominance reversal (Figure 6A). This is because limited dispersal, through an excess of homozygosity, diminishes the effects of dominance on both: (i) the expected frequency *p*_0_ at which the deleterious allele is maintained before environmental change; and (ii) selection when beneficial. In fact, as dispersal becomes increasingly limited, the trajectory profiles of alleles that experience a dominance shift become almost indistinguishable from the profiles of alleles that did not (Figure 6B).

**Figure 6:**
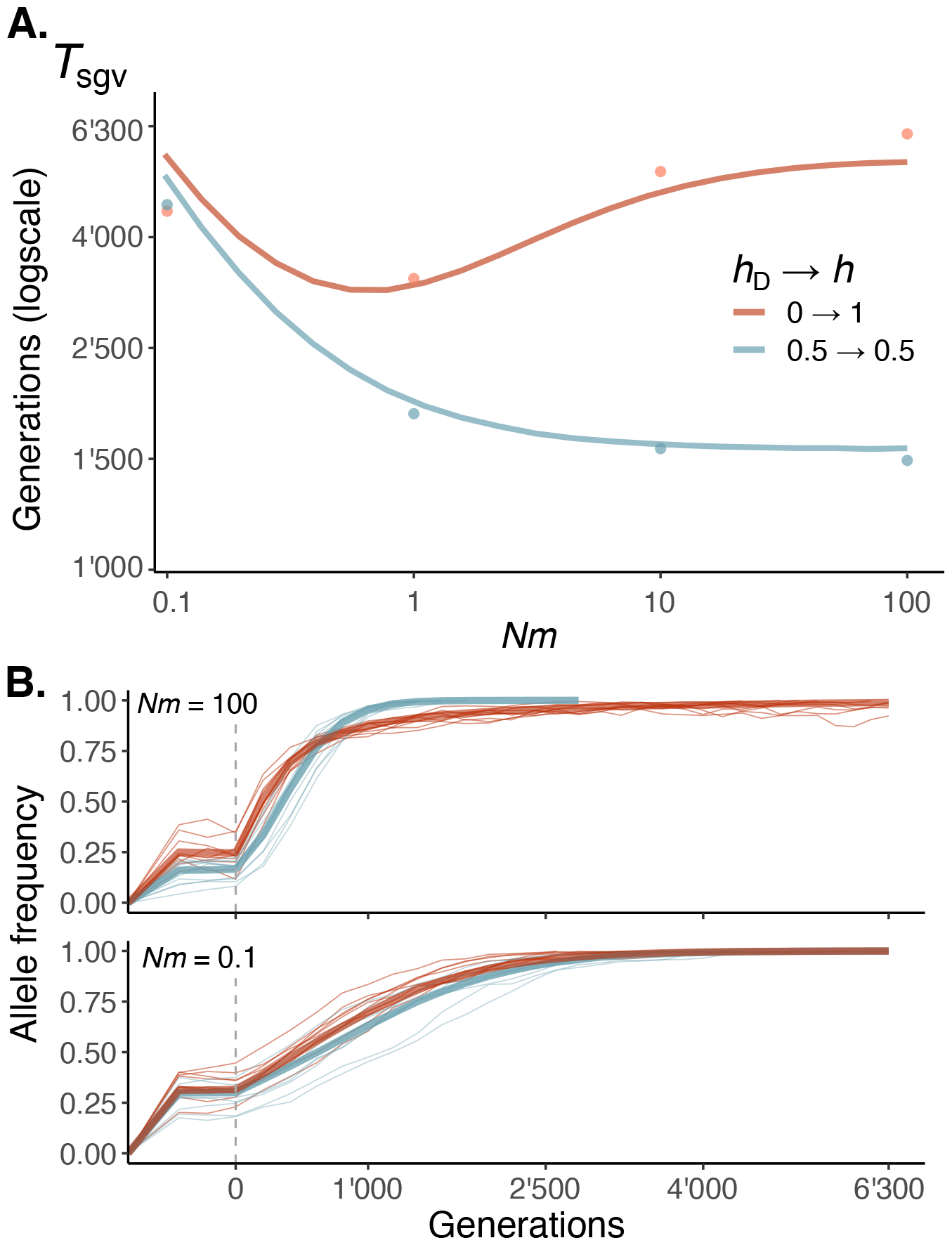
The effects of dominance reversal under limited dispersal. **A**. Expected time *T*_sgv_ of fixation (on a log scale) of recessive deleterious alleles that become beneficial dominant (*h*_D_ = 0 and *h =* 1, in red), and of additive alleles (*h*_D_ *= h =* 0.5 in blue) with solid lines from diffusion approximation (eq. 13 with eqs. 7 and A49), and dots for the average from individual based simulations (300 replicates for each set of parameters, section B in File S1 for details). Parameters: same as Figure 5. **B**. Fixation trajectories of alleles showing dominance reversal (*h*_D_ = 0 and *h =* 1, in red) and additive alleles (*h*_D_ *= h =* 0.5 in blue) in a well-mixed (top, *Nm =* 100) and dispersal-limited (bottom, *Nm =* 0.1) population. Environmental change takes place at *t =* 0 (dashed vertical line). For each scenario (in different colours, see A for legend), thin lines show ten randomly sampled trajectories, and thick lines show the mean trajectory among all trajectories. Parameters: same as A.

### 3.4 Longer waiting but faster fixation under extinction-recolonization

We have assumed that demes are of fixed and constant size. But deme extinctions, whereby entire local populations vanish and their habitat is made available for recolonization, are ecologically and evolutionarily relevant as they modulate the consequences of dispersal (Pannell and Charlesworth, 1999; Rousset, 2004). To explore how the interplay between extinction-recolonization and limited dispersal influences fixation times, we assume that before step (i) of the life cycle (see section 2.1), each deme independently goes extinct with a probability 0 ≤ *e* < 1, in which case all individuals present in that deme die before producing any gamete (section C in File S1 for details). Each extinct deme is then available for recolonization by 2*N* gametes from extant demes during dispersal. We sample these 2*N* gametes in two ways to examine contrasting scenarios of recolonization (as in Slatkin, 1977; Whitlock and McCauley, 1990): (i) in the propagule model, gametes are sampled from the gametic pool of a single extant deme, which is chosen at random among all extant demes; while (ii) in the migrant pool model, gametes are sampled from the joint gametic pool of all extant demes.

We look at the total time of taken for a *de novo* beneficial mutation, *T*_new_ (eq. 12), which depends on the waiting time for a fixing allele to arise and on the time to fixation (Figure 7A). We find that deme extinctions tend to have limited effects on *T*_new_, unless the mode of recolonization follows the propagule model and dispersal is strongly limited (third row of Figure 7A). In this case, *T*_new_ is greater under deme extinctions mostly due to an inflation in waiting time. This is because by increasing the covariance in allele frequency among demes, extinction-recolonization dynamics reduces effective population size relative to census size (Figure 7B; Slatkin, 1977; Whitlock and McCauley, 1990; Barton, 1993; Barton and Whitlock, 1997). This reduction is especially significant under the propagule model because in this case, a recolonized deme and the deme of origin for the propagule have the same allele frequency on average, which boosts the covariance among demes (Figure 7B, bottom). The resulting increase in genetic drift reduces the fixation probability of beneficial alleles, and thus in turn, increases the waiting time (first term of eq. 12; Figure 7C, bottom). The increase in genetic drift also causes a reduction in fixation time (second term of eq. 12), but this does not compensate for the inflated waiting time in the case of propagule recolonization (Figure 7A). Altogether, our results indicate that adaptation from *de novo* mutations is characterized by faster fixations separated by longer waiting times under extinction-recolonization dynamics.

**Figure 7:**
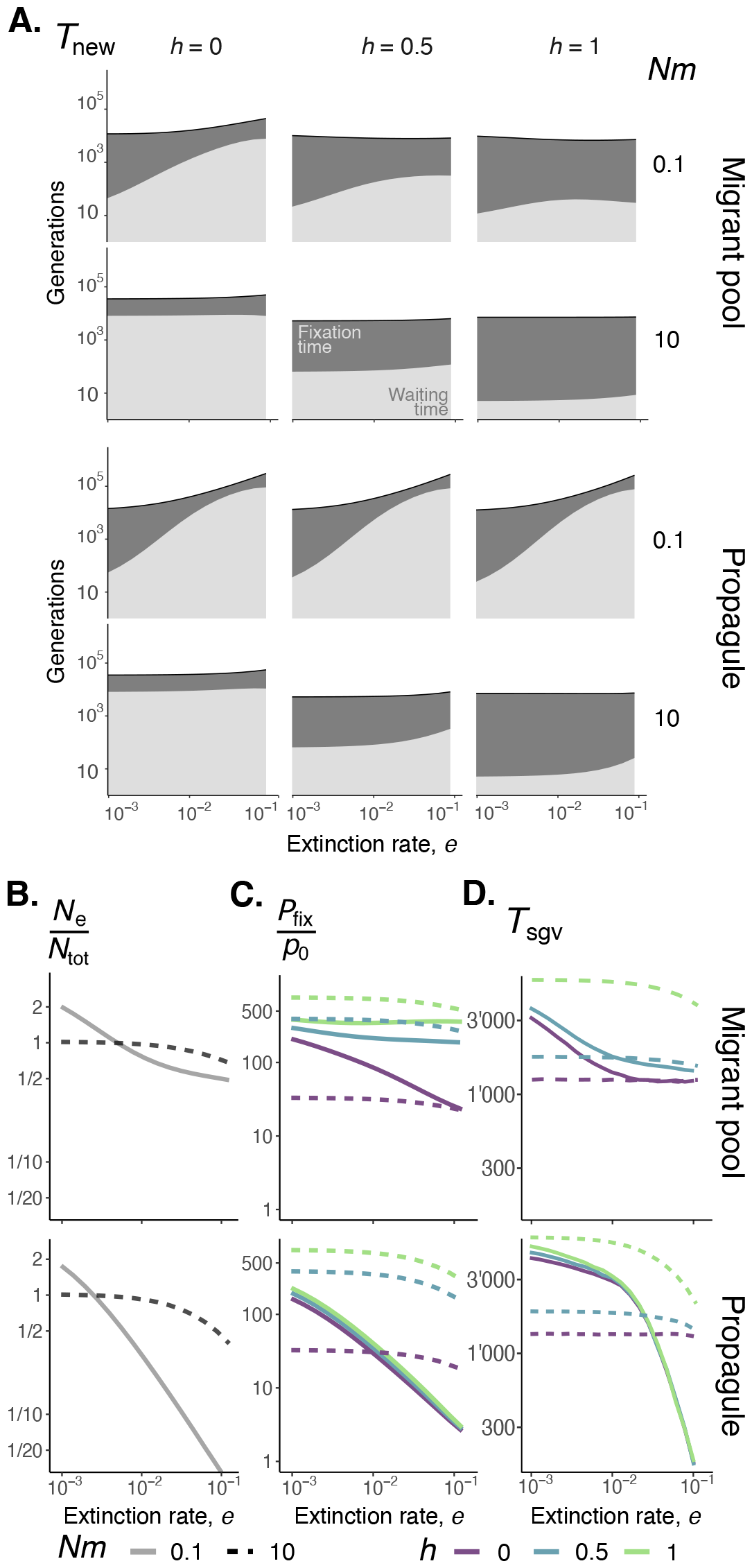
The effects of local extinctions and recolonization dynamics. **A**. Expected total time *T*_new_ of fixation (on a log scale) of recessive (*h =* 0, left), additive (*h =* 0.5, middle) and dominant (*h =* 1, right) *de novo* mutations, *p*_0_ = 1/(2*N*_T_), for different dispersal rates (*Nm =* 10 in bottom, and 0.1 in top) and recolonization models (migrant pool model in top row and propagule model in bottom, also for panels B-D), from eq. (12) (with eqs. 6 and 7). Parameters: same as Figure 1. **B**. Effective population size relative to census size, *N*_e_/*N*_T_ (from eq. C4 in File S1). Dashed line for *Nm =* 10 and full line for *Nm =* 0.1 (also for panels C-D). Parameters: same as A. **C**. Fixation probabilities normalised to initial frequency *p*_0_ of recessive (*h =* 0),additive (*h =* 0 .5) and dominant (*h =* 1) alleles (in different colours, see legend) arising as single copies *p*_0_ = 1/ (2*N*_T_), from eq. (6) with *Nm =* 10 (dashed) and 0.1 (full), and under different recolonization models (top and bottom rows). Parameters: same as A. **D**. Expected time *T*_sgv_ of fixation (on a log scale) of standing recessive (*h =* 0), additive (*h =* 1/2) and dominant (*h =* 1) variants (in different colors) from eq. 13 (with eqs. 7 and A49). The case *h =* 1 with *Nm =* 0.1 is omitted as comparisons with simulations showed a poor fit (Figure C1 in File S1). Parameters: same as Figure 5.

In contrast to fixation of *de novo* mutations, extinctions almost always reduce the expected number of generations a beneficial allele from standing genetic variation takes to fix, *T*_sgv_ (eq. 13, Figure 7D). This is because the waiting time is no longer relevant when the allele is already present in the population. In fact, the reduction in *N*_e_ owing to extinction-recolonization dynamics accelerates adaptation as it both: (i) reduces fixation time; and (ii) leads to on average a greater frequency *p*_0_ of *A* at the time of environmental change. As a result, fixation of standing genetic variants can be significantly quicker when extinctions are common and recolonization follows the propagule model (Figure 7D, bottom).

## 4 Discussion

Our analyses indicate that limited dispersal can accelerate the fixation of beneficial *de novo* alleles when: (i) dispersal is mildly limited; and (ii) allelic effects on fitness are not too weak and are far from additive (e.g. *h* < 0.1 or *h* > 0.9 in Figure 1B). This may be relevant to natural populations as the dispersal rates under which we found that recessive and dominant mutations fix quicker than under panmixia lead to *F*_ST_ levels that agree with estimates from a wide range of taxa (roughly *Nm* > 1 so on average 1 or more migrants per generation, leading to *F*_ST_ < 0.2, Figure 1A and B; e.g. fish, Ståhl, 1981; Glover et al., 2013; crustaceans, Benzie, 2000; plants, Giles and Goudet, 1997; Potenko and Velikov, 1998; Tamaki et al., 2008; insects, Irvin et al., 1998; Kumar and Singh, 2017; birds, Forstmeier et al., 2007; pp. 302-303 in Hartl and Clark, 2007 for an overview). Additionally, the notion that alleles have nonadditive fitness effects is supported by multiple lines of evidence, both theoretical (Fisher, 1928; Wright, 1934; Kacser and Burns, 1981; Manna et al., 2011; Billiard et al., 2021) and empirical, with mutations thought to be often at least partially recessive, with an average dominance coefficient *h* close to 0.2 (Mukai et al., 1972; Agrawal and Whitlock, 2011; Huber et al., 2018; reviewed in Orr, 2010 and Li and Bank, 2023). Further, the selection coefficient that is required to observe a decrease in the time to fixation under limited dispersal (*N*_T_*s* greater than 50) sits well within empirically estimated fitness effects (Eyre-Walker and Keightley, 2007). We considered specifically the case where *N*_T_*s =* 200 in our main text figures, which corresponds to a 1% increase in fecundity due to a single substitution in *N*_d_ = 200 demes of *N =* 100 individuals (as in Roze and Rousset, 2003). We tested the effect of stronger selection with simulations whose results are shown in Figure B in File S1. These show similar patterns to our baseline model, i.e. mild dispersal limitation speeds up fixation when alleles are recessive or dominant. In fact, strong selection tends to amplify this effect (Figure B in File S1).

In addition to the time to fixation, the pace of adaptation also depends on the waiting time for a fixing mutation to appear (eq. 12; Glémin and Ronfort, 2013). Because limited dispersal reduces most significantly the waiting time for a fixing recessive allele to appear, the total time for a *de novo* mutation to fix is most shortened when beneficial alleles are recessive (purple region in Figure 4B). Overall, our results thus suggest that with all else being equal, a subdivided population should be better adapted and show greater mean fecundity than a panmictic population, provided dispersal is only mildly limited and adaptation is driven by recessive *de novo* mutations.

In contrast, limited dispersal always slows down fixation of standing genetic variants that are recessive before and after they become beneficial due to an environmental change (Figure 5A, purple line). Rather, mild dispersal limitation tends to accelerate the fixation of dominant alleles here (Figure 5A, green line). This is because limited dispersal leads to a greater boost in frequency of a deleterious allele when it is dominant compared to when it is recessive (Figure 5B, compare top to bottom). Nevertheless, the time taken for dominant genetic variants to fix in response to changes in selective pressures is typically greater than recessive variants, although limited dispersal tends to reduce this difference (Figure 5A).

More broadly, limited dispersal diminishes the importance of genetic dominance on the time taken by alleles to fix. This is in part because limited dispersal leads to inbreeding, which causes an excess of homozygotes whose fecundity does not depend on genetic dominance. Models involving partial selfing (or assortative mating), which also causes elevated homozygosity, similarly found lesser importance of dominance for fixation (Roze and Rousset, 2004; Glémin and Ronfort, 2013; Newberry et al., 2016; Hartfield and Bataillon, 2020; Charlesworth, 2020). Our model, however, contrasts with these scenarios because limited dispersal also: (i) leads to kin competition, which reduces the overall strength of selection; and (ii) increases effective population size *N*_e_ whereas selfing alone reduces *N*_e_. These two effects explain why strongly limited dispersal always delays fixation, whereas selfing generally speeds up fixation (Roze and Rousset, 2004; Glémin and Ronfort, 2013; Newberry et al., 2016; Hartfield and Bataillon, 2020; Charlesworth, 2020). In fact, our results under extinction-recolonization dynamics align more closely with those under selfing as limited dispersal reduces *N*_e_ when extinctions are sufficiently common (Figure 7B).

Through its effects on time to fixation, limited dispersal may have implications for the signature of selective sweeps. The idea behind this is that when a selected allele goes to fixation more rapidly, there are fewer opportunities for recombination so that genetic diversity at nearby neutral sites tends to be reduced, leading to what is referred to as a *hard sweep*; whereas when fixation is slow, recombination is more likely to break the association between an adaptive allele and its original background before fixation, leading to a *soft sweep* (Hermisson and Pennings, 2005, 2017; Messer and Petrov, 2013; Jensen, 2014). More specifically, the linkage disequilibrium between a new beneficial allele and a linked neutral allele decreases at a rate given by their recombination rate *r* in a large well-mixed population, i.e. linkage disequilibrium decays as exp(−*r t*), where *t* is the number of generations that the fixing allele takes to rise to high frequency (Maynard Smith and Haigh, 1974; Barton, 2000). Accordingly, the probability of observing a hard sweep is lower under limited dispersal if limited dispersal increases *t* (Barton, 2000; Pennings and Hermisson, 2006d; Kim and Maruki, 2011). However, modelling studies have found contrasting effects of limited dispersal on the signature of sweeps, which is typically quantified by *F*_ST_ at linked neutral loci. In fact, the fixation of a beneficial allele can increase or decrease *F*_ST_, depending on initial conditions and on dominance (Slatkin and Wiehe, 1998; Santiago and Caballero, 2005; Kim and Maruki, 2011; Teshima and Przeworski, 2006; Roze and Rousset, 2008; Ewing et al., 2011). In particular, *F*_ST_ at linked neutral loci is expected to increase when a recessive allele sweeps, whereas *F*_ST_ is expected to decrease when a dominant allele sweeps (eq. 79 in Roze and Rousset, 2008). We performed simulations of evolution at two linked loci where one is neutral and initially polymorphic, and the other is under positive selection (section D in File S1 for details). The results we find align with those of Roze and Rousset (2008). When beneficial alleles are additive, dispersal has no effect on the probability of observing a hard sweep, i.e. on the probability that the polymorphism at the neutral locus is lost with fixation of the beneficial allele (blue line in panel A, Figure C in File S1). This is because although the time to fixation is greater, and so are recombination opportunities under limited dispersal, most of the new haplotypes created by recombination are lost due to local drift within demes (panel B, Figure C in File S1). Meanwhile, limited dispersal increases the probability of observing a hard sweep for a recessive allele and decreases it for a dominant allele so that these probabilities converge to that of an additive allele (purple and green lines in panel A, Figure C in File S1). This is because limited dispersal decreases (respectively, increases) the proportion of time that a recessive (dominant) allele spends at low frequency (Figure 2), thus affecting the recombination opportunities with new backgrounds for these alleles.

The association between a selected allele and its original background can also be broken when recurrent mutations create beneficial mutations that are identical-by-state and that fix with different backgrounds (Pennings and Hermisson, 2006c; Ralph and Coop, 2010; Paulose et al., 2019). How likely this is to happen can be inferred from comparing the waiting and fixation time (eq. 12). If the fixation time is long compared to waiting time, then recurrent mutations should be more likely to lead to a soft sweep. Inspection of Figure 4 reveals that the time to fixation can become longer than the waiting time as dispersal becomes more limited, especially if mutations are common. This suggests that limited dispersal may favour soft sweeps through recurrent mutations. To investigate more definitively how limited dispersal affects the signature of sweeps, it would be interesting to extend our model to consider multiple linked loci (e.g., extending Roze and Rousset, 2008 to finite number of demes or Lehmann and Rousset, 2009 to limited dispersal).

Finally, our results are based on several assumptions. First, we assumed that dispersal is gametic, which is relevant for plant and marine taxa but less so for terrestrial animals where it is often zygotes that disperse. But provided that mating is random within demes and demes are large enough, alellic segregation is similar under gametic and zygotic dispersal (Roze and Rousset, 2003). Second, we assumed that selection is soft, i.e. that each deme produces the same number of gametes. We explore the case of hard selection in section E in File S1 such that demes showing greater frequency of allele *A* produce more gametes. Hard selection reduces by a small margin the time to fixation, but does not affect our results otherwise (Figure E1 in File S1). Third, the diffusion approximation also relies on the assumption that demes are homogeneous and that dispersal is uniform among them (i.e. no isolation-by-distance). Isolation by distance in principle delays fixation (Rousset, 2006), unless demes show specific patterns of connectivity that create sources and sinks that may facilitate fixation (e.g. Marrec et al., 2021). Fourth, we focused on the expectation of the number of generations taken for fixation, which may be misleading if the underlying distribution is fat tailed and skewed towards large values. To check for this, we computed the median time to fixation using individual based simulations. We found that the mean and the median are close, indicating that the distribution of times to fixation is fairly symmetrical around the mean (Figure D in File S1).

## Acknowledgements

We thank Ehouarn Le Faou, Claudia Bank, Denis Roze and Michael Whitlock for discussions, Nick Barton for useful feedback on our manuscript, and the Swiss National Science Foundation for funding (PCEFP3181243 to CM).

## Data Availability Statement

The code used for our simulations is available here: https://vsudbrack.github.io/LimitedDispersalSM. Otherwise, the authors affirm that all data necessary for confirming the conclusions of the article are present within the article and figures.

