## Supplementary File S1 for "Fixation times of *de novo* and standing beneficial variants in subdivided populations"

### Supplementary Material

### File S1

for

### Fixation times of *de novo* and standing beneficial variants in subdivided populations

Vitor Sudbrack and Charles Mullon [1]

#### List of supplementary texts

A Diffusion approximation for baseline model

B Computer simulations

C Extinction-recolonization dynamics

D Hard and soft sweeps

E Hard selection

#### List of supplementary figures

Figure A Selection threshold for faster fixation under dispersal limitation

Figure B Fixation under strong selection

Figure C Effects of dispersal limitation on the signature of sweeps

Figure D Summary statistics of fixation events

### A Diffusion approximation for baseline model

Here, we briefly describe how we used the diffusion approximation to model the segregation of allele $A$ following Roze and Rousset (2003) for the island model of dispersal (see also Barton, 1993; Whitlock, 2003; Cherry and Wakeley, 2003; Cherry, 2003; Wakeley, 2003; Wakeley and Takahashi, 2004). Readers familiar with this approach may skip this supplementary text, which we present for the sake of completeness.

The diffusion approximation relies on two quantities: the expectation of, and the variance in, the change in allele frequency over one generation (Crow and Kimura, 1970; Ewens, 2004). To specify this frequency change, consider a given generation $t$. Let $p_{ij1} \in \{0,1\}$ and $p_{ij2} \in \{0,1\}$ be the frequency of $A$ at each homologous chromosomes of individual $j \in \{1,\ldots,N\}$ in deme $i \in \{1,\ldots,N_\mathrm{d}\}$ at this generation $t$. The allele frequency then is

$$p_{ij} = \frac{p_{ij1} + p_{ij2}}{2} \tag{A1}$$

in that individual, and

$$p = \frac{1}{N_\mathrm{T}} \sum_{i=1}^{N_\mathrm{d}} \sum_{j=1}^{N} p_{ij} \tag{A2}$$

in the whole population (recall $N_\mathrm{T} = N \cdot N_\mathrm{d}$). We use a prime symbol to denote quantities in the next generation $t+1$, so that $p'$ refers to the frequency of $A$ in the next generation, and thus

$$\Delta p = p' - p \tag{A3}$$

is the allele frequency change.

#### A.1 Expected allele frequency change

Here, we derive eq. (1) of the main text, i.e. the expected frequency change over one generation, $\mathrm{E}[\Delta p | p]$, to the first order of a small parameter $\delta$ such that $s \sim \mathcal{O}(\delta)$ and $N_\mathrm{d} \sim \mathcal{O}(1/\delta)$. Under random chromosomal segregation and in the absence of mutation, this expectation can be written as

$$\mathrm{E}[\Delta p | p] = \mathrm{E}\left[ \underbrace{\frac{1}{2N_\mathrm{T}} \sum_{i=1}^{N_\mathrm{d}} \sum_{j=1}^{N} \mathrm{E}[w_{ij} p_{ij} | \mathbf{P}] - p}_{\mathrm{E}[\Delta p | \mathbf{P}]} \;\middle|\; p \right], \tag{A4}$$

where $\mathbf{P}$ is the set of realised genotypes of all individuals, i.e. $\mathbf{P} = \left\{p_{ijk} : k \in \{1,2\}, j \in \{1,\ldots,N\}, i \in \{1,\ldots,N_\text{d}\}\right\}$, and $w_{ij}$ in eq. (A4) is the expected number of successful gametes (i.e. that are recruited) produced by individual $j$ from deme $i$, given the genotype of every individual in its generation. We refer to $w_{ij}$ as the fitness of individual $j$ from deme $i$.

##### A.1.1 Fitness

To specify the fitness $w_{ij}$ of a focal individual, we first connect relative fecundity $z_{ij}$ of this individual with its genotype through the expression

$$z_{ij} = 1 + s\left[2hp_{ij} + (1-2h)p_{ij1}p_{ij2}\right] , \tag{A5}$$

such that $aa$, $aA$, and $AA$ individuals have relative fecundity 1, $1+hs$, and $1+s$, respectively (as per the model described in section 2.1 of the main text; eq. 11 in Roze and Rousset, 2003 with their $z_\text{aa} = 1$). In the limit where absolute fecundity is large, fitness for our baseline model can be written as

$$w_{ij} = \underbrace{2N\cdot(1-m)\cdot\frac{z_{ij}}{\sum_{k=1}^{N} z_{ik}}}_{\text{philopatric fitness}} + \underbrace{\sum_{\substack{r=1\\ r\neq i}}^{N_\text{d}} 2N\cdot\frac{m}{N_\text{d}-1}\cdot\frac{z_{ij}}{\sum_{k=1}^{N} z_{ik}}}_{\text{dispersal fitness}} , \tag{A6}$$

which consists in the sum of philopatric fitness (expected number of gametes recruited in their deme of origin) and dispersal fitness (recruited in all other demes). Equation (A6) simplifies to

$$w_{ij} = 2\cdot\frac{z_{ij}}{z_i}, \tag{A7}$$

where

$$z_i = \frac{1}{N}\sum_{j=1}^{N} z_{ij} \tag{A8}$$

is the average fecundity in deme $i$.

More generally, we consider in the next two sections models where individual fitness can be written to the first order of $\delta$ as a function of three variables,

$$w_{ij} = w(z_{ij}, z_i, z) + \mathcal{O}\left(\delta^2\right), \tag{A9}$$

where

$$z = \frac{1}{N_\text{d}}\sum_{i=1}^{N_\text{d}} z_i \tag{A10}$$

is the population average fecundity. One useful property of fitness is that since the total population size $N_{\mathrm{T}}$ is constant, the average population fitness is also fixed at $\sum_{i=1}^{N_{\mathrm{d}}}\sum_{j=1}^{N} w_{ij}/N_{\mathrm{T}} = 2$. As a consequence, we also have

$$\left(\frac{\partial w_{ij}}{\partial z_{ij}} + \frac{\partial w_{ij}}{\partial z_i} + \frac{\partial w_{ij}}{\partial z}\right)_{s=0} = 0 \tag{A11}$$

(p. 96 in Rousset, 2004).

##### A.1.2 Conditional expected allele frequency change

To compute eq. (A4), we follow Roze and Rousset (2003) and first characterise the expected allele frequency change conditioned on all genotypes, $\mathrm{E}[\Delta p|\mathbf{P}]$. Plugging eq. (A9) into eq. (A4) and performing a Taylor expansion around $s = 0$ obtains

$$\mathrm{E}[\Delta p|\mathbf{P}] = \frac{s}{2N_{\mathrm{T}}}\sum_{i=1}^{N_{\mathrm{d}}}\sum_{j=1}^{N}\left(\frac{\partial w_{ij}}{\partial z_{ij}}\frac{dz_{ij}}{ds} + \frac{\partial w_{ij}}{\partial z_i}\frac{dz_i}{ds} + \frac{\partial w_{ij}}{\partial z}\frac{dz}{ds}\right)p_{ij} + \mathcal{O}\left(\delta^2\right), \tag{A12}$$

where here and hereafter all derivatives are estimated under neutrality (i.e., $s = \delta = 0$ and thereby $z_{ij} = z_i = z = 1$). Using eqs. (A5), (A8) and (A10), eq. (A12) then becomes

$$\begin{aligned}\mathrm{E}[\Delta p|\mathbf{P}] = \frac{s}{2N_{\mathrm{T}}}\sum_{i=1}^{N_{\mathrm{d}}}\sum_{j=1}^{N}\Bigg(&\frac{\partial w_{ij}}{\partial z_{ij}}\left[2hp_{ij} + (1-2h)p_{ij1}p_{ij2}\right]\\ &+\frac{1}{N}\sum_{k=1}^{N}\frac{\partial w_{ij}}{\partial z_i}\left[2hp_{ik} + (1-2h)p_{ik1}p_{ik2}\right]\\ &+\frac{1}{N_{\mathrm{T}}}\sum_{r=1}^{N_{\mathrm{d}}}\sum_{k=1}^{N}\frac{\partial w_{ij}}{\partial z}\left[2hp_{rk} + (1-2h)p_{rk1}p_{rk2}\right]\Bigg)p_{ij} + \mathcal{O}\left(\delta^2\right).\end{aligned} \tag{A13}$$

This can be written after some straightforward rearrangements as

$$\begin{aligned}\mathrm{E}[\Delta p|\mathbf{P}] = hs&\left(\frac{\partial w_{ij}}{\partial z_{ij}}\left\langle p_{ij}^2\right\rangle_{ij} + \frac{\partial w_{ij}}{\partial z_i}\left\langle p_i^2\right\rangle_i + \frac{\partial w_{ij}}{\partial z}p^2\right)\\ &+(1-2h)\frac{s}{2}\left(\frac{\partial w_{ij}}{\partial z_{ij}}\left\langle p_{ij1}p_{ij2}\right\rangle_{ij} + \frac{\partial w_{ij}}{\partial z_i}\left\langle p_{ij1}p_{ij2}p_i\right\rangle_{ij} + \frac{\partial w_{ij}}{\partial z}p\left\langle p_{ij1}p_{ij2}\right\rangle_{ij}\right)\\ &+\mathcal{O}\left(\delta^2\right),\end{aligned} \tag{A14}$$

where we used the convention

$$\left\langle f_{ij}\right\rangle_{ij} = \frac{1}{N_{\mathrm{T}}}\sum_{i=1}^{N_{\mathrm{d}}}\sum_{j=1}^{N} f_{ij}, \qquad \left\langle f_{ij}\right\rangle_j = \frac{1}{N}\sum_{j=1}^{N} f_{ij}, \qquad \left\langle f_i\right\rangle_i = \frac{1}{N_{\mathrm{d}}}\sum_{i=1}^{N_{\mathrm{d}}} f_i, \tag{A15}$$

for any quantity $f_{ij}$ that can be ascribed to an individual $ij$, and the property that

$$\langle p_{ij1}p_{ij2}p_{ij}\rangle_{ij} = \langle p_{ij1}p_{ij2}\rangle_{ij} \tag{A16}$$

since $p_{ij1}$ and $p_{ij2}$ are indicator variables. Equation (A14) is equivalent to eq. (14) in Roze and Rousset (2003). Using eq. (A11) to substitute for $\frac{\partial w_{ij}}{\partial z}$, eq. (A14) becomes

$$\begin{aligned}\mathrm{E}[\Delta p|\mathbf{P}] = hs&\left[\frac{\partial w_{ij}}{\partial z_{ij}}\left(\left\langle p_{ij}^2\right\rangle_{ij} - p^2\right) + \frac{\partial w_{ij}}{\partial z_i}\left(\langle p_i^2\rangle_i - p^2\right)\right]\\ &+ (1-2h)\frac{s}{2}\left[(1-p)\frac{\partial w_{ij}}{\partial z_{ij}}\langle p_{ij1}p_{ij2}\rangle_{ij} + \frac{\partial w_{ij}}{\partial z_i}\langle p_{ij1}p_{ij2}(p_i - p)\rangle_{ij}\right] + \mathcal{O}(\delta^2)\ .\end{aligned} \tag{A17}$$

##### A.1.3 Unconditional expected allele frequency change

We then plug eq. (A17) into eq. (A4) to obtain

$$\begin{aligned}\mathrm{E}[\Delta p|p] = hs&\left(\frac{\partial w_{ij}}{\partial z_{ij}}\left(\mathrm{E}\left[\left\langle p_{ij}^2\right\rangle_{ij}\middle|p\right] - p^2\right) + \frac{\partial w_{ij}}{\partial z_i}\left(\mathrm{E}\left[\langle p_i^2\rangle_i\middle|p\right] - p^2\right)\right)\\ &+ (1-2h)\frac{s}{2}\left((1-p)\frac{\partial w_{ij}}{\partial z_{ij}}\mathrm{E}\left[\langle p_{ij1}p_{ij2}\rangle_{ij}\middle|p\right] + \frac{\partial w_{ij}}{\partial z_i}\mathrm{E}\left[\langle p_{ij1}p_{ij2}(p_i-p)\rangle_{ij}\middle|p\right]\right)\\ &+ \mathcal{O}(\delta^2),\end{aligned} \tag{A18}$$

which depends on various moments of the genetic distribution in the population. These only need to be evaluated under neutrality as they are already weighed by $s$ in eq. (A18). In that case, they can be connected to probabilities of identity-by-descent under neutrality (Roze and Rousset, 2003, for more details). In fact,

$$\mathrm{E}\left[\langle p_{ij1}p_{ij2}\rangle_{ij}\middle|p\right] = r_0^{\mathrm{D}}p + \left(1-r_0^{\mathrm{D}}\right)p^2 + \mathcal{O}(\delta) \tag{A19}$$

$$\mathrm{E}\left[\left\langle p_{ij}^2\right\rangle_{ij}\middle|p\right] = r_0^{\mathrm{R}}p + \left(1-r_0^{\mathrm{R}}\right)p^2 + \mathcal{O}(\delta) \tag{A20}$$

$$\mathrm{E}\left[\langle p_i^2\rangle_i\middle|p\right] = r_1^{\mathrm{R}}p + \left(1-r_1^{\mathrm{R}}\right)p^2 + \mathcal{O}(\delta) \tag{A21}$$

$$\mathrm{E}\left[\langle p_{ij1}p_{ij2}p_i\rangle_{ij}\middle|p\right] = a^{\mathrm{R}}p + \left(1-a^{\mathrm{R}}-c^{\mathrm{R}}\right)p^2 + c^{\mathrm{R}}p^3 + \mathcal{O}(\delta) \tag{A22}$$

where $r_0^{\mathrm{D}}$ is the probability that the two homologous genes of the same individual are identical-by-descent (IBD);

$$r_0^{\mathrm{R}} = \frac{1}{2} + \frac{1}{2}r_0^{\mathrm{D}} \tag{A23}$$

is the probability that two genes sampled with replacement in the same individual are IBD; $r_1^{\mathrm{R}}$ is the probability that two genes sampled from the same deme with replacement are IBD; $a^{\mathrm{R}}$ is the probability that two homologous genes of an individual coalesce with a third gene (sampled from the same deme at random), and

$$c^{\mathrm{R}} = 1 - a^{\mathrm{R}} - \left(r_0^{\mathrm{D}} - a^{\mathrm{R}}\right) - 2\left(r_1^{\mathrm{R}} - a^{\mathrm{R}}\right) \tag{A24}$$

is the probability that none of these three genes are IBD (see eq. A-23 in Flintham et al., 2021 for details on eq. A24).

Plugging eqs. (A19) through (A22) into eq. (A18) yields

$$\begin{aligned}\mathrm{E}[\Delta p|p] = hs\left(\frac{\partial w_{ij}}{\partial z_{ij}} r_0^{\mathrm{R}} + \frac{\partial w_{ij}}{\partial z_i} r_1^{\mathrm{R}}\right) p(1-p) \\ + (1-2h)\,\frac{s}{2}\left(\frac{\partial w_{ij}}{\partial z_{ij}}\left[r_0^{\mathrm{D}} + (1-r_0^{\mathrm{D}})p\right] + \frac{\partial w_{ij}}{\partial z_i}\left[a^{\mathrm{R}} + 2(r_1^{\mathrm{R}} - a^{\mathrm{R}})p\right]\right) p(1-p) + \mathcal{O}\left(\delta^2\right),\end{aligned} \tag{A25}$$

which is equivalent to eq. (23) in Roze and Rousset (2003) with our eq. (A24) substituted into it. We compute probabilities of identity-by-descent in section A.3 in terms of dispersal and deme size.

From eq. (A7), we have

$$\frac{\partial w_{ij}}{\partial z_{ij}} = 2 \qquad \text{and} \qquad \frac{\partial w_{ij}}{\partial z_i} = -2\,, \tag{A26}$$

which substituted into eq. (A25) and after some re-arrangements using eq. (A23) yields eq. (1) of the main text.

#### A.2 Variance in frequency change

Here, we derive eqs. (3) and (4) of the main text, i.e. the variance in frequency change over one generation, $\mathrm{V}[\Delta p|p]$, to the leading order of $\delta$. Following eq. A8 in Roze and Rousset (2003), we decompose this change as

$$\mathrm{V}[\Delta p|p] = \frac{1}{{N_{\mathrm{d}}}^2}\sum_{i=1}^{N_{\mathrm{d}}} \mathrm{V}\left[p_i'|p\right] + \frac{1}{{N_{\mathrm{d}}}^2}\sum_{i=1}^{N_{\mathrm{d}}}\sum_{\substack{r=1\\ r\neq i}}^{N_{\mathrm{d}}} \mathrm{C}\left[p_i', p_r'\middle|p\right]\,, \tag{A27}$$

where recall $p_i' = \sum_{j=1}^{N} p_{ij}' \big/ N$ is the frequency of $A$ in deme $i$ at generation $t+1$. Since sampling of gametes occurs independently in each deme in our baseline model, the covariance term vanishes, leaving us with

$$\mathrm{V}[\Delta p|p] = \frac{1}{{N_{\mathrm{d}}}^2}\sum_{i=1}^{N_{\mathrm{d}}} \mathrm{V}\left[p_i'|p\right] = \frac{1}{N_{\mathrm{d}}}\left\langle \mathrm{V}\left[p_i'|p\right]\right\rangle_i \tag{A28}$$

as in eq. (A1) in Roze and Rousset (2003). Since $\left\langle \mathrm{V}\left[p_i'|p\right]\right\rangle_i$ is multiplied by $1/N_\mathrm{d}$ in eq. (A28), we can compute it assuming the number of demes is infinite and there is no selection ($s = 0$). To do so, we take a slight detour from the derivation of Roze and Rousset (2003) (though we eventually get the same result). Let us first condition on all genetic information $\mathbf{P}$ at the previous generation (in particular, in knowing all deme frequencies) using the law of total variance,

$$\mathrm{V}[\Delta p|p] = \frac{1}{N_\mathrm{d}}\left\langle \mathrm{E}\left[\mathrm{V}[p_i'|\mathbf{P}]\Big|p\right] + \mathrm{V}\left[\mathrm{E}[p_i'|\mathbf{P}]\Big|p\right]\right\rangle_i. \tag{A29}$$

To characterise this, let $N_{\mathrm{M},i}$ be the realized number of migrant gametes received in deme $i$ at generation $t+1$ from all the other demes. Under neutrality, $N_{\mathrm{M},i}$ is distributed binomially with parameters,

$$N_{\mathrm{M},i} \overset{\mathrm{law}}{\sim} \mathrm{Binom}(2N, m)\ . \tag{A30}$$

Conditional on $N_{\mathrm{M},i}$, the allele frequency at focal deme $i$ has mean and variance given by,

$$\begin{aligned}\mathrm{E}\left[p_i'|\mathbf{P}, N_{\mathrm{M},i}\right] &= \left(\frac{2N - N_{\mathrm{M},i}}{2N}\right)p_i + \left(\frac{N_{\mathrm{M},i}}{2N}\right)p + \mathcal{O}(\delta) & \text{(A31)}\\ \mathrm{V}\left[p_i'|\mathbf{P}, N_{\mathrm{M},i}\right] &= \frac{N_{\mathrm{M},i}}{(2N)^2}p(1-p) + \frac{2N - N_{\mathrm{M},i}}{(2N)^2}p_i(1-p_i) + \mathcal{O}(\delta)\,. & \text{(A32)}\end{aligned}$$

Marginalizing those over $N_{\mathrm{M},i}$ (eq. A30), we obtain

$$\begin{aligned}\mathrm{E}\left[p_i'|\mathbf{P}\right] &= (1-m)p_i + m\,p + \mathcal{O}(\delta) & \text{(A33)}\\ \mathrm{V}\left[p_i'|\mathbf{P}\right] &= \mathrm{V}\left[\mathrm{E}\left[p_i'|\mathbf{P}, N_{\mathrm{M},i}\right]|\mathbf{P}\right] + \mathrm{E}\left[\mathrm{V}\left[p_i'|\mathbf{P}, N_{\mathrm{M},i}\right]|\mathbf{P}\right] & \\ &= \frac{m(1-m)(p_i-p)^2 + mp(1-p) + (1-m)p_i(1-p_i)}{2N} + \mathcal{O}(\delta)\,. & \text{(A34)}\end{aligned}$$

Plugging eqs. (A33) and (A34) into eq. (A29), one obtains

$$\mathrm{V}[\Delta p|p] = \frac{1}{N_\mathrm{d}}\left(\frac{p - [1-(1-m)^2]p^2 - (1-m)^2\left\langle \mathrm{E}\left[{p_i}^2|p\right]\right\rangle_i}{2N} + (1-m)^2\left\langle \mathrm{V}[p_i|p]\right\rangle_i\right) + \mathcal{O}\left(\delta^2\right) \tag{A35}$$

after re-arrangements and using the fact that $\mathrm{E}\left[p_i|p\right] = p$.

We can simplify eq. (A35) further by noticing first that $\mathrm{V}[\langle p_i\rangle_i|p] = \langle \mathrm{V}[p_i|p]\rangle_i$ since the covariance between deme frequencies is zero (when the number of demes is infinite). We also have that by definition, $\mathrm{V}[\langle p_i\rangle_i|p] = \mathrm{V}[p|p] = 0$. These relationships then entail that $\langle \mathrm{V}[p_i|p]\rangle_i = 0$. Second, from eq. (A21) we have

$$\left\langle \mathrm{E}\left[{p_i}^2|p\right]\right\rangle_i = \mathrm{E}\left[\left\langle {p_i}^2\right\rangle_i|p\right] = r_1^\mathrm{R}p + \left(1 - r_1^\mathrm{R}\right)p^2 + \mathcal{O}(\delta)\ . \tag{A36}$$

Substituting for these into eq. (A35) yields,

$$\mathrm{V}[\Delta p|p] = p(1-p)\frac{1-(1-m)^2 r_1^{\mathrm{R}}}{2N_{\mathrm{d}}N} + \mathcal{O}\left(\delta^2\right) \tag{A37}$$

which is equal to eq. (A4) of Roze and Rousset (2003) using the fact that $r_1^{\mathrm{D}} = (1-m)^2 r_1^{\mathrm{R}}$.

Following standard theory, we define the (variance) effective population size $N_{\mathrm{e}}$ such that

$$\mathrm{V}[\Delta p|p] = \frac{p(1-p)}{2N_{\mathrm{e}}} + \mathcal{O}\left(\delta^2\right), \tag{A38}$$

which by comparison to eq. (A37) leads to

$$N_{\mathrm{e}} = \frac{N_{\mathrm{d}}N}{1-(1-m)^2 r_1^{\mathrm{R}}} \tag{A39}$$

and using $r_1^{\mathrm{D}} = (1-m)^2 r_1^{\mathrm{R}}$ obtains eq. (4) of the main text. Note also that because $r_1^{\mathrm{D}} = F_{\mathrm{ST}}$ in the island model of dispersal, eq. (4) is equal to eq. (9.48) in Rousset (2004). We compute $r_1^{\mathrm{D}}$ in eq. (A44). Plugging for this into eq. (4) then gives

$$N_{\mathrm{e}} = \left(1 + \frac{1}{2N}\cdot\frac{(1-m)^2}{1-(1-m)^2}\right)N_{\mathrm{d}}N \tag{A40}$$

in terms of the dispersal rate $m$.

#### A.3 Probabilities of coalescence

Here, we briefly explain how relevant probabilities of IBD are derived using standard coalescent arguments (e.g. Karlin, 1968; Wang, 1997; Rousset, 2004). Since these probabilities already weigh on terms of order $\delta$ in the expected and variance in allelic frequency change, they are derived under the assumption that there is no selection and the number of demes is effectively infinite. This latter assumption means in particular that the probability that two genes sampled in different demes coalesce is zero.

##### A.3.1 Pairwise coalescence probability

Recall that $r_0^{\mathrm{D}}$ denotes the probability of coalescence between the two homologous genes of a focal individual, $r_0^{\mathrm{R}}$ the probability of coalescence between two genes sampled with replacement in the same individual (eq. A23), and $r_1^{\mathrm{R}}$ the probability of coalescence between two genes sampled with replacement from the same deme. These last two probabilities are connected by

$$r_1^{\mathrm{R}} = \frac{1}{N} r_0^{\mathrm{R}} + \left(1 - \frac{1}{N}\right) r_1^{\mathrm{D}} \tag{A41}$$

where $r_1^{\mathrm{D}}$ is the probability of coalescence of two different genes sampled in the same deme (superscripts R and D generally denote samplings of genes with and without replacement, respectively, while subscripts 0 and 1 denote samplings of genes within the same individual and within the same deme, respectively). The first term of eq. (A41) corresponds to the probability of sampling the same individual, and the second, of sampling different individuals.

The probability of coalescence of two different genes in a given deme after one generation can be expressed as

$$r_1^{\mathrm{D}\prime} = (1-m)^2 \Bigg[ \underbrace{\frac{1}{N}\left(\frac{1}{2} + \frac{1}{2} r_0^{\mathrm{D}}\right)}_{\text{(i)}} + \underbrace{\left(1 - \frac{1}{N}\right) r_1^{\mathrm{D}}}_{\text{(ii)}} \Bigg], \tag{A42}$$

where $(1-m)^2$ is the probability of sampling two philopatric genes, and we have decomposed their coalescence as to whether they have the same parent (i) or not (ii). However, since we assume that gametes fuse randomly in each deme following dispersal, we have

$$r_0^{\mathrm{D}} = r_1^{\mathrm{D}}, \tag{A43}$$

or equivalently, $F_{\mathrm{IT}} = F_{\mathrm{ST}}$ (and $F_{\mathrm{IS}} = 0$). Substituting for $r_0^{\mathrm{D}}$ into eq. (A42) and solving the recursion for $r_1^{\mathrm{D}\prime} = r_1^{\mathrm{D}}$ yields

$$r_0^{\mathrm{D}} = r_1^{\mathrm{D}} = \frac{(1-m)^2}{2N - (1-m)^2(2N-1)}. \tag{A44}$$

This together with eqs. (A23) and (A41) allows us to compute $r_1^{\mathrm{R}}$, which gives all the required components for the selection gradient when $h = 1/2$ (eq. 1). Equation (A44) is equivalent to eq. (3.14) for $F_{\mathrm{ST}}$ in Rousset (2004) considering diploids (i.e., their $N$ as $2N$).

##### A.3.2 Threeway coalescence probability

When $h \neq 1/2$, the selection gradient further depends on the probability $a^{\mathrm{R}}$ that two homologous genes of an individual coalesce with a third gene sampled with replacement from the same deme. This probability can be expressed as

$$a^{\mathrm{R}} = \frac{1}{N} r_0^{\mathrm{D}} + \left(1 - \frac{1}{N}\right) a^{\mathrm{D}}, \tag{A45}$$

where $a^{\mathrm{D}}$ is the probability that two homologous genes of an individual coalesce with a third gene of a different individual from the same deme (i.e. sampled without replacement). This probability satisfies the recursion,

$$a^{\mathrm{D}\prime} = \underbrace{(1-m)^3}_{\substack{\text{three philopatric}\\ \text{genes}}} \left[ \underbrace{\left(\frac{1}{2N}\right)^2}_{\text{(i)}} + \underbrace{3\frac{1}{2N}\left(1-\frac{1}{2N}\right) r_1^{\mathrm{D}}}_{\text{(ii)}} + \underbrace{\left(1-\frac{1}{2N}\right)\left(1-\frac{2}{2N}\right) a^{\mathrm{D}}}_{\text{(iii)}} \right] \tag{A46}$$

where (i) captures immediate coalescence of these three copies in the same gene; (ii) captures immediate coalescence of only two alleles, followed by coalescence with the third gene further in the past with probability $r_1^{\mathrm{D}}$; and finally, (iii) captures the case where none of the three genes coalesce in the previous generation. Solving eq. (A46) for $a^{\mathrm{D}\prime} = a^{\mathrm{D}}$ yields

$$a^{\mathrm{D}} = \frac{\left[1+3\left(2N-1\right)r_1^{\mathrm{D}}\right](1-m)^3}{\left(2N\right)^2 - \left(2N-1\right)\left(2N-2\right)(1-m)^3}\ , \tag{A47}$$

which substituted into eq. (A45) with eq. (A44) gives $a^{\mathrm{R}}$ as required for the selection gradient (eq. 1).

#### A.4 Stationary frequency distribution

In section 3.3 of the main text, we consider the expected number of generations taken by alleles to fix whose initial frequencies are sampled from a stationary frequency distribution $\phi(p)$. Here, we detail how we computed this distribution $\phi(p)$. According to the assumptions of this model (section 3.3 for details), the expected change in frequency is given by

$$\begin{aligned} \mathrm{E}[\Delta p|p] = \mu(1-2p) - s_{\mathrm{D}}p(1-p)\Big[ & p + r_0^{\mathrm{D}}(1-p) + h_{\mathrm{D}}(1-r_0^{\mathrm{D}})(1-2p) \\ & - \left(r_1^{\mathrm{R}} + (r_1^{\mathrm{R}} - a^{\mathrm{R}})(2h_{\mathrm{D}}-1)(1-2p)\right)\Big] + \mathcal{O}\left(\delta^2\right)\ , \end{aligned} \tag{A48}$$

where the first term captures mutation with $\mu \sim \mathcal{O}(\delta)$, and the second negative selection with parameters $s_{\mathrm{D}}$ and $h_{\mathrm{D}}$. The variance in allele frequency change is given by eq. (A38) with eq. (A40). We can then compute the stationary allele frequency distribution under the diffusion approximation by solving the Fokker-Planck equation with eqs. (A48) and (A38) (Crow and Kimura, 1970, p. 372), such that

$$\phi(p) = \frac{c_0}{p(1-p)} \exp\left(2\int_0^p \gamma(x)dx\right)\ , \tag{A49}$$

where $c_0$ is a constant that guarantees $\int_0^1 \phi(p)dp = 1$, and $\gamma(x)$ is given by eq. (5) with eqs. (A48), (3) and (4) plugged into it (eq. A49 is found in Crow and Kimura, 1970, p. 425; note we incorporated their factor $2N_{\mathrm{e}}$ into $c_0$).

### B Computer simulations

#### B.1 Basic algorithm

The Python code used for our simulations is available here:

https://vsudbrack.github.io/LimitedDispersalSM

These simulations track the number of adults at each deme $i = 1, \dots, N_\mathrm{d}$ according to their genotypes: $n_i^{aa}, n_i^{aA}, n_i^{AA}$ (with $n_i^{aa} + n_i^{aA} + n_i^{AA} = N$) over time according to the following algorithm.

Given $n_i^{aa}, n_i^{aA}, n_i^{AA}$, we first calculate the proportion of $A$ gametes produced in each deme $i$ with

$$p_i^{(\mathrm{S})} = \underbrace{\frac{1}{2N + 2hsn_i^{aA} + 2sn_i^{AA}}}_{\text{total number of gametes}} \underbrace{\left[(1+hs)n_i^{aA} + 2(1+s)n_i^{AA}\right]}_{\text{total A gametes}}, \tag{B1}$$

where the superscript (S) is for selection. Second, mutations occur, so that this proportion becomes,

$$p_i^{(\mathrm{M})} = p_i^{(\mathrm{S})} + \underbrace{\mu\left(1 - p_i^{(\mathrm{S})}\right)}_{a \to A} - \underbrace{\mu\, p_i^{(\mathrm{S})}}_{A \to a}, \tag{B2}$$

in each deme $i$. Third, dispersal alters the proportion of $A$ gametes in each deme $i$ to,

$$p_i^{(\mathrm{D})} = (1-m)p_i^{(\mathrm{M})} + m\, p_{(-i)}^{(\mathrm{M})}, \tag{B3}$$

where $p_{(-i)}^{(\mathrm{M})} = \frac{N_\mathrm{d}}{N_\mathrm{d}-1} p^{(\mathrm{M})} - \frac{1}{N_\mathrm{d}-1} p_i^{(\mathrm{M})}$ (with $p^{(\mathrm{M})}$ as the mean across all demes) is the average of $p_j^{(\mathrm{M})}$ among all demes excluding $i$. Finally, gametes fuse randomly within demes and density-dependent regulation occurs, so that the number of adults of each genotype after one iteration of the life cycle is thus given by

$$\left[n_i^{aa}, n_i^{aA}, n_i^{AA}\right]' \overset{\text{law}}{\sim} M_3\left(N;\ \left(1-p_i^{(\mathrm{D})}\right)^2,\ 2p_i^{(\mathrm{D})}\left(1-p_i^{(\mathrm{D})}\right),\ \left(p_i^{(\mathrm{D})}\right)^2\right), \tag{B4}$$

where $M_3$ is a Multinomial distribution with three classes.

#### B.2 Analyses

##### B.2.1 Time to fixation of *de novo* mutations

To study the time to fixation of *de novo* mutations (conditioned on fixation), we iterate the life cycle starting with all $aa$ adults but one heterozygous $aA$. We let 40'000 simulations run with no mutation

– i.e., $\mu = 0$ in eq. (B2) so that eventually, $A$ is fixed or lost. We recorded the number of generations until fixation whenever $A$ fixed.

###### B.2.2 Fixation of standing genetic variants

To study fixation of standing genetic variants, we first let the population evolve for 10'000 generations under selection, mutation and drift. We inspected that simulations reached equilibrium, which they did very rapidly (e.g. Figure 5C). During this time, mutation rate is set to $N_{\mathrm{T}}\mu = 2$, selection and dominance coefficients in eq. (B1) is set to $s = -s_{\mathrm{D}} = -0.001$ and $h = h_{\mathrm{D}}$ (see Figures for values). At $t = 10'001$, selection on $A$ becomes positive $s = 0.01 > 0$ and to study the time to fixation from standing variation, we then set $\mu = 0$. We record the number of generations until the eventual fixation of allele $A$.

### C Extinction-recolonization dynamics

Here we describe our analyses to investigate the impact of extinction-recolonization dynamics, whose results are described in section 3.4. We also present the algorithm to simulate extinction and recolonization dynamics.

#### C.1 Expected allele frequency change

When selection is soft, fitness eq. (A7) still holds with extinction-recolonization occurring as described in section 3.4 (eq. 35 of Roze and Rousset, 2003 for rationale; section E.3 here for the case of hard selection). Consequently, the expected change in allele frequency is still given by eq. (1), i.e. by

$$\begin{aligned}\mathrm{E}[\Delta p|p] = sp(1-p)\Big[p + r_0^{\mathrm{D}}(1-p) + h(1-r_0^{\mathrm{D}})(1-2p) \\ - \left(r_1^{\mathrm{R}} + (r_1^{\mathrm{R}} - a^{\mathrm{R}})(2h-1)(1-2p)\right)\Big] + \mathcal{O}\left(\delta^2\right).\end{aligned} \tag{C1}$$

What does change due to extinction-recolonization are the probabilities of coalescence that appear in this equation. We derive those below (eqs. C6 and C8).

#### C.2 Variance in allele frequency change

Due to recolonization, the covariance in gene frequency between demes can no longer be ignored in eq. (A27). As shown by Roze and Rousset (2003), the variance in allele frequency change can still be written as

$$\mathrm{V}[\Delta p|p] = \frac{p(1-p)}{2N_{\mathrm{e}}} + \mathcal{O}\left(\delta^2\right), \tag{C2}$$

but now the effective population size is given by

$$\begin{aligned} N_{\mathrm{e}} &= \left(\frac{1-e}{2r_1^{\mathrm{R}}\left[1-(1-e)^2(1-m)^2\right]}\right)N_{\mathrm{d}} && \text{(C3)}\\ &= \left(\frac{1-e}{2\left(r_0^{\mathrm{R}} + (N-1)r_1^{\mathrm{D}}\right)\left[1-(1-e)^2(1-m)^2\right]}\right)N_{\mathrm{d}}N && \text{(C4)}\end{aligned}$$

(eq. 29 in Roze and Rousset, 2003; then using eq. A41). We compute $r_1^{\mathrm{R}}$ for our extinction-recolonization model below.

#### C.3 Probabilities of coalescence under local extinctions

##### C.3.1 Pairwise coalescence probability

After one generation, the probability of coalescence of two alleles sampled without replacement from the same deme can be expressed as

$$r_1^{\mathrm{D}'} = (1-e)\underbrace{(1-m)^2\left[\frac{1}{N}\left(\frac{1}{2}+\frac{1}{2}r_0^{\mathrm{D}}\right)+\left(1-\frac{1}{N}\right)r_1^{\mathrm{D}}\right]}_{\text{no extinction, eq. (A42)}}+e\underbrace{\varphi\left[\frac{1}{N}\left(\frac{1}{2}+\frac{1}{2}r_0^{\mathrm{D}}\right)+\left(1-\frac{1}{N}\right)r_1^{\mathrm{D}}\right]}_{\text{extinction}}, \qquad \text{(C5)}$$

where $\varphi$ is the probability that two randomly sampled gametes among the $2N$ colonizers come from the same deme. We model recolonization in two ways: (i) with a migrant pool model, in which each gamete comes from different demes so $\varphi = 0$; and (ii) with a propagule model, wherein all gametes come from the same randomly sampled deme so $\varphi = 1$. As the labels indicate, the first term of eq. (C5) captures coalescence in the case where the deme did not go extinct, and the second term where it did. Equation (C5) differs, although only marginally, to eq. (B20) of Roze and Rousset (2003) because they assume two rounds of reproduction within recolonization dynamics. As before, $r_0^{\mathrm{D}} = r_1^{\mathrm{D}}$. Substituting for this into eq. (C5) and solving for $r_1^{\mathrm{D}'} = r_1^{\mathrm{D}}$, yields,

$$r_0^{\mathrm{D}} = r_1^{\mathrm{D}} = \frac{(1-e)(1-m)^2+e\varphi}{2N-[(1-e)(1-m)^2+e\varphi](2N-1)}\,. \qquad \text{(C6)}$$

The probability $r_1^{\mathrm{R}}$ is found by plugging eq. (C6) into eq. (A41).

##### C.3.2 Threeway coalescence probability

The probability $a^{\mathrm{D}}$ that the two homologous genes of an individual plus a third gene randomly sampled from the remaining genes in that same deme, are all IBD is given by solving the recursion

$$a^{\mathrm{D}'} = \left((1-e)(1-m)^3+e\varphi^2\right)\left[\left(\frac{1}{2N}\right)^2+3\frac{1}{2N}\left(1-\frac{1}{2N}\right)r_1^{\mathrm{D}}+\left(1-\frac{1}{2N}\right)\left(1-\frac{2}{2N}\right)a^{\mathrm{D}}\right]. \qquad \text{(C7)}$$

This is similar in structure but slightly different to eq. (B21) of Roze and Rousset (2003) owing to our different assumptions about recolonization. Solving eq. (C7) for $a^{\mathrm{D}'} = a^{\mathrm{D}}$ yields,

$$a^{\mathrm{D}} = \frac{\left(1+3(2N-1)r_1^{\mathrm{D}}\right)\left[(1-m)^3(1-e)+e\varphi^2\right]}{(2N)^2-(2N-1)(2N-2)\left[(1-m)^3(1-e)+e\varphi^2\right]}, \qquad \text{(C8)}$$

where $r_1^{\mathrm{D}}$ is given by eq. (C6).

##### C.4 Simulations

To simulate evolution with extinctions and recolonization dynamics, we extend our basic algorithm described in section B.1. Before the production of gametes, we sample a random number $n_\mathrm{e}$ of demes that go extinct from a binomial distribution with parameters,

$$n_\mathrm{e} \overset{\text{law}}{\sim} \mathrm{Binom}(N_\mathrm{d}, e). \tag{C9}$$

Those $n_\mathrm{e}$ demes are sampled uniformly without replacement from all demes.

Extinct demes do not produce any gametes and therefore do not participate in the dispersal phase (i.e., $p_{(-i)}^{(\mathrm{M})}$ in eq. B3 is averaged only over extant demes).

Extinct demes are available for recolonization and this can occur in two ways depending on the model. In the migrant pool model of recolonization, each deme $i$ among these $n_\mathrm{e}$ demes is recolonized by sampling gametes from a multinomial distribution

$$\left[n_i^{aa}, n_i^{aA}, n_i^{AA}\right]' \overset{\text{law}}{\sim} M_3\left(N;\ \left(1-p_\mathrm{e}\right)^2,\ 2p_\mathrm{e}\left(1-p_\mathrm{e}\right),\ \left(p_\mathrm{e}\right)^2\right), \tag{C10}$$

wherein $p_\mathrm{e}$ is the average of gamete frequencies $p_i^{(\mathrm{M})}$ before dispersal among all $N_\mathrm{d}-n_\mathrm{e}$ extant demes.

In the propagule model, one deme $r$ among all $N_\mathrm{d}-n_\mathrm{e}$ extant demes is chosen at random to recolonize a given extinct deme $i$. The multinomial sampling to establish the adults in the next adult generation is

$$\left[n_i^{aa}, n_i^{aA}, n_i^{AA}\right]' \overset{\text{law}}{\sim} M_3\left(N;\ \left(1-p_r^{(\mathrm{M})}\right)^2,\ 2p_r^{(\mathrm{M})}\left(1-p_r^{(\mathrm{M})}\right),\ \left(p_r^{(\mathrm{M})}\right)^2\right). \tag{C11}$$

Comparisons between these simulations and predictions from the diffusion approximation reveal that that the diffusion approximation typically works well under extinction-recolonization dynamics (Figure C1), except with propagule recolonization and very weak dispersal, $Nm = 0.1$. In this case, the diffusion approximation tends to overestimate the time taken by alleles to fix, especially if the allele is dominant and the extinction rate is high.

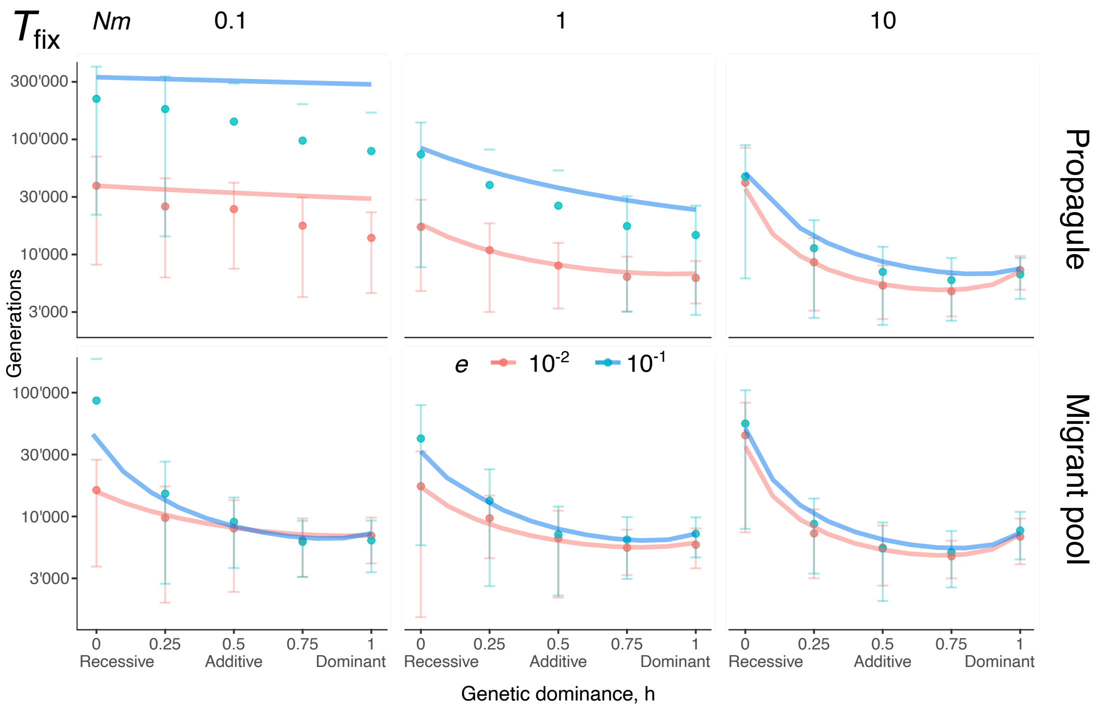


**Figure C1: Comparison with simulations under local extinctions**. Expected time to fixation $T_{\text{fix}}(p_0)$ (on a log scale) of alleles arising as single copies $p_0 = 1/(2N_{\text{T}})$ when $e = 10^{-1}$ and $10^{-2}$, for different dispersal rates (columns) and models of recolonization (rows). Solid lines are the numerical integration of eq. (7), and dots are the average of simulations (180 replicates for each set of parameters, error bars show standard deviation). Parameters: same as Figure 1.

### D Hard and soft sweeps

To investigate the probability of hard and soft sweeps (and produce Figure C here), we simulated evolution at two linked loci, one under selection and another neutral. Each locus is diallelic with, as before, an adapted allele $A$ and a wild-type allele $a$ at the locus under selection, and neutral alleles $B$ and $b$ at the neutral locus.

The algorithm has the same broad structure as in section B.1, with modifications to accommodate two linked loci. Briefly, we track the number of adults $n_i^{X|Y}$ that carry genotypes made up of haplotypes $X$ and $Y$ (with $X, Y \in \{AB, Ab, aB, ab\}$) in each deme $i$. Given these numbers, we can straightforwardly calculate gametic frequencies after selection and recombination in each deme $i$ ($p_{X,i}^{(S)}$). For instance, the frequency of gametes with haplotype $AB$ before dispersal is

$$p_{AB,i}^{(S)} \propto (1+s)n_i^{AB|AB} + \frac{1}{2}(1+s)n_i^{AB|Ab} + \frac{1}{2}(1+hs)(1-r)n_i^{AB|ab} + \frac{1}{2}(1+hs)rn_i^{Ab|aB}, \qquad \text{(D1)}$$

where $r$ is the recombination rate and the proportionality is such that gametic frequencies sum to one (i.e. such that $p_{AB,i}^{(S)} + p_{Ab,i}^{(S)} + p_{aB,i}^{(S)} + p_{ab,i}^{(S)} = 1$). Gametic dispersal is implemented the same way as in eq. (B3) for each of the four gametic types. Finally, gametes fuse randomly within each deme and density-dependent regulation occurs. This means the number of adults of each genotype is given by a Multinomial sampling, with probabilities built from the product of relevant gametic frequencies after dispersal (i.e., similar to eq. B4).

Initially, haplotypes are either $aB$ or $ab$ with equal probability, so that wild-type $a$ is fixed and, neutral $B$ and $b$ are equally frequent on average. To reflect how dispersal and drift shaped variation among demes before the appearance of beneficial allele $A$, we use eq. 2 of Cherry and Wakeley, 2003 (with their $\overline{x} = 0.5$) and sample the frequency in each deme $i$ of haplotype $aB$ from an appropriate Beta distribution (the frequency of haplotype $ab$ is given by the reciprocal of the frequency of $aB$). We form the first generation of adults from these frequencies (i.e. obtain $n_i^{X|Y}$ for each $X, Y$) by assuming Hardy-Weinberg equilibrium within each deme. One individual of this first generation is chosen at random in the whole population and mutates at its selected locus, from $a$ to $A$.

We let the population evolve until the adaptive allele $A$ is either fixed or lost. Among all the simulations where $A$ fixes, we record the frequency of $B$ at the neutral locus, and refer to the sweep as a *hard sweep* if either $B$ or $b$ is also fixed at the time $A$ fixes. Otherwise, we refer to the sweep as a *soft sweep.*

For each set of parameters, we perform 10'000 replicates and calculate the proportion $\pi_{\text{HS}}$ of hard sweeps among those replicates where $A$ fixes. Results are shown in Figure C here.

### E Hard selection

As a baseline, we have assumed that all demes produce an equal number of gametes, i.e. that selection is soft. Here, we consider the case of hard selection, whereby demes in which the frequency of $A$ is greater produce more gametes. We find overall that fixations occur marginally faster under hard selection than under soft selection, but otherwise our results under hard selection are qualitatively similar to those under soft selection.

#### E.1 Expected and variance in allele frequency change

Under hard selection, the fitness of an individual $j$ in deme $i$ can be written as

$$w(z_{ij}, z_i, z) = \underbrace{\frac{2(1-m)z_{ij}}{(1-m)z_i + mz}}_{\text{philopatric fitness}} + \underbrace{\frac{2mz_{ij}}{z}}_{\text{dispersal fitness}}, \tag{E1}$$

which yields

$$\frac{\partial w_{ij}}{\partial z_{ij}} = 2 \qquad \text{and} \qquad \frac{\partial w_{ij}}{\partial z_i} = -2(1-m)^2. \tag{E2}$$

(evaluated at $s = 0$, i.e. $z_{ij} = z_i = z = 1$; ). Plugging these derivatives into eq. (A25), we obtain that the expected change in allele frequency is

$$\begin{aligned}\mathrm{E}[\Delta p|p] = & sp(1-p)\Big[p + r_0^{\mathrm{D}}(1-p) + h(1-r_0^{\mathrm{D}})(1-2p) \\ & - (1-m)^2\Big(r_1^{\mathrm{R}} + (r_1^{\mathrm{R}} - a^{\mathrm{R}})(2h-1)(1-2p)\Big)\Big] + \mathcal{O}(\delta^2).\end{aligned} \tag{E3}$$

Comparing eq. (E3) with the case of soft selection (eq. 1) reveals that under hard selection, kin competition is reduced by $(1-m)^2$. This means that effective selection is stronger under hard selection.

The relevant probabilities of coalescence are not affected by whether selection is soft or hard since we only need these probabilities under neutrality. We can thus use those computed in sections A.3 (eqs. A44 and A47). This also entails that the variance in allele frequency change is still written as

$$\mathrm{V}[\Delta p|p] = \frac{p(1-p)}{2N_{\mathrm{e}}} + \mathcal{O}(\delta^2), \tag{E4}$$

with the effective population size given by eq. (A40), i.e.

$$N_{\mathrm{e}} = \left(1 + \frac{1}{2N} \cdot \frac{(1-m)^2}{1-(1-m)^2}\right) N_{\mathrm{d}} N. \tag{E5}$$

#### E.2 Analyses

We used the above to compute the total time to fixation of *de novo* mutations (i.e. the sum of waiting and fixation times, eq. 12 with eqs. 6 and 7, Figure E1A) and the time to fixation of standing genetic variants (eq. 13 with eqs. 7 and A49) where dominance is preserved (Figure E1B) and reversed (Figure E1C). We focused on the case where demes are small, $N = 10$, because this is where differences among hard and soft selection are the greatest (Roze and Rousset, 2003). Figure E1 shows that hard selection speeds up adaptation regardless if it stems from new *de novo* mutations or standing genetic variation. This is because kin competition is weaker and so effective selection is stronger under hard selection. The quantitative consequences of this for fixation times, however, are small (compare dashed and solid lines in Figure E1).

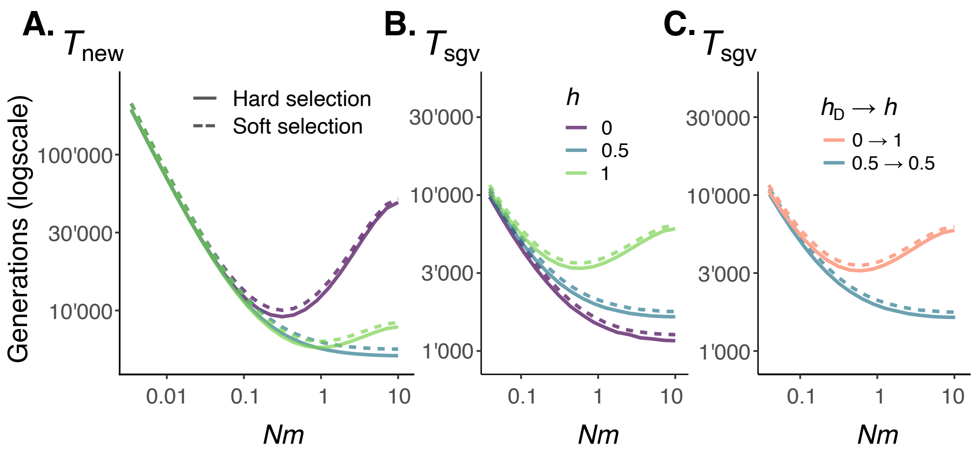


**Figure E1: Fixation times under hard selection. A.** Expected total time to fixation of *de novo* mutants $T_{new}$ (on a log scale) that are recessive ($h = 0$), additive ($h = 0.5$) and dominant ($h = 1$, in different colors) with $p_0 = 1/(2N_T)$ under hard (full line) and soft (dashed line) selection. Parameters: $N_d = 2000$, $N = 10$, $N_T\mu = 0.02$, $s = 0.01$, so that $N_T$ is the same as in the main text. **B.** Expected time of fixation of standing genetic variants $T_{sgv}$ (on a log scale) with $h_D = h$ for recessive ($h = 0$), additive ($h = 0.5$) and dominant ($h = 1$) variants under hard (full line) and soft (dashed line) selection. Parameters: $s_D = 10^{-3}$, $N_T\mu = 2$, other parameters same as A. **C.** Expected time of fixation of standing genetic variants $T_{sgv}$ (on a log scale) of recessive deleterious variant that becomes beneficial dominant ($h_D = 0$ and $h = 1$, orange), and an additive allele ($h_D = h = 1/2$, blue) under hard (full line) and soft (dashed line) selection. Parameters: same as B.

#### E.3 Hard selection and local extinction

We also investigated the case of hard selection under extinction-recolonization dynamics. In that case, fitness is given by,

$$w(z_{ij}, z_i, z) = (1-e)\underbrace{\left[\frac{2(1-m)z_{ij}}{(1-m)z_i + mz} + \frac{2mz_{ij}}{z}\right]}_{\text{as in eq. (E1) if not extinct}} + \underbrace{(1-e)\cdot 2N(eN_d)\cdot\frac{z_i}{(1-e)N_d z}\cdot\frac{z_{ij}}{Nz_i}}_{\text{descendants via recolonization}}, \tag{E6}$$

(as in eq. 38 of Roze and Rousset, 2003) such that

$$\frac{\partial w_{ij}}{\partial z_{ij}} = 2 \qquad \text{and} \qquad \frac{\partial w_{ij}}{\partial z_i} = -2(1-e)(1-m)^2\,, \tag{E7}$$

which plugged into eq. (A25) gives

$$\begin{aligned}\mathrm{E}[\Delta p|p] = sp(1-p)\Big[&p + r_0^{\mathrm{D}}(1-p) + h(1-r_0^{\mathrm{D}})(1-2p)\\ &-(1-e)(1-m)^2\Big(r_1^{\mathrm{R}} + (r_1^{\mathrm{R}} - a^{\mathrm{R}})(2h-1)(1-2p)\Big)\Big] + \mathcal{O}\big(\delta^2\big).\end{aligned} \tag{E8}$$

Note that fitness is not influenced as to whether recolonization occurs according to the migrant pool or propagule model. Probabilities of coalescence are not influenced by hard selection and we thus use those computed in section C.3. Effective population size, meanwhile, is given by eq. (C4). Our results for time of fixation of *de novo* mutations and standing genetic variants are shown in Figure E2. This shows that hard selection accelerates adaptation, especially under the propagule model of recolonization where dispersal is weak (Figure E2A and B, bottom left).

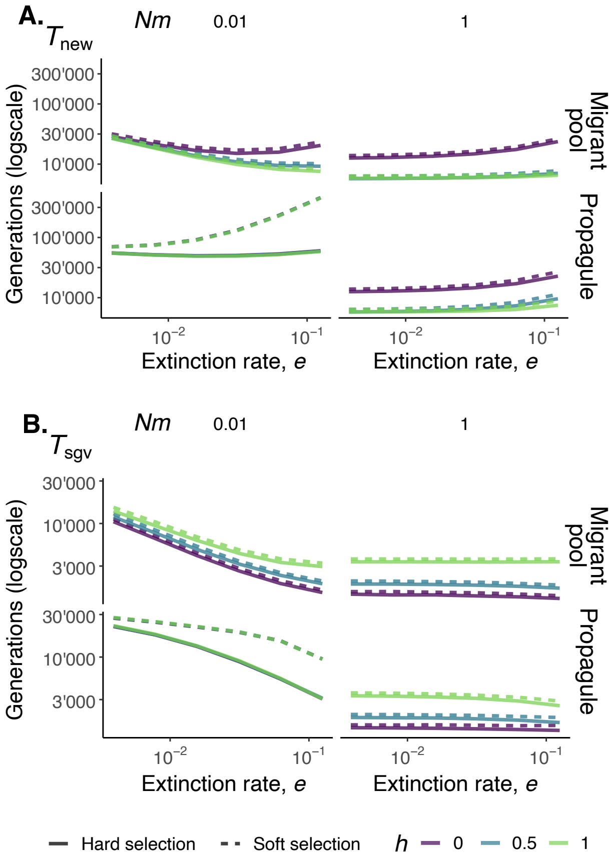


**Figure E2: The effects of extinction and recolonization dynamics under hard selection**. **A.** Expected total time of fixation of *de novo* mutations $T_{\text{new}}$ (on a log scale) with recessive ($h = 0$), additive ($h = 0.5$) and dominant ($h = 1$) effects (in different colors) arising as single copies $p_0 = 1/(2N_T)$ under hard (full line) and soft (dashed line) selection for $Nm = 0.01$ (left) and $Nm = 1$ (right), and under the migrant pool (top) and propagule (bottom) model of recolonization. Other parameters: same as Figure E1A. **B.** Expected time of fixation of standing genetic variants $T_{\text{sgv}}$ (on a log scale) with $h_D = h$ for recessive ($h =$ 0), additive ($h = 0.5$) and dominant ($h = 1$) alleles under hard (full line) and soft (dashed line) selection for $Nm = 0.1$ (left) and $Nm = 1$ (right), and under the migrant pool (top) and propagule (bottom) model of recolonization. $h_D = h$. Parameters: same as Figure E1B.

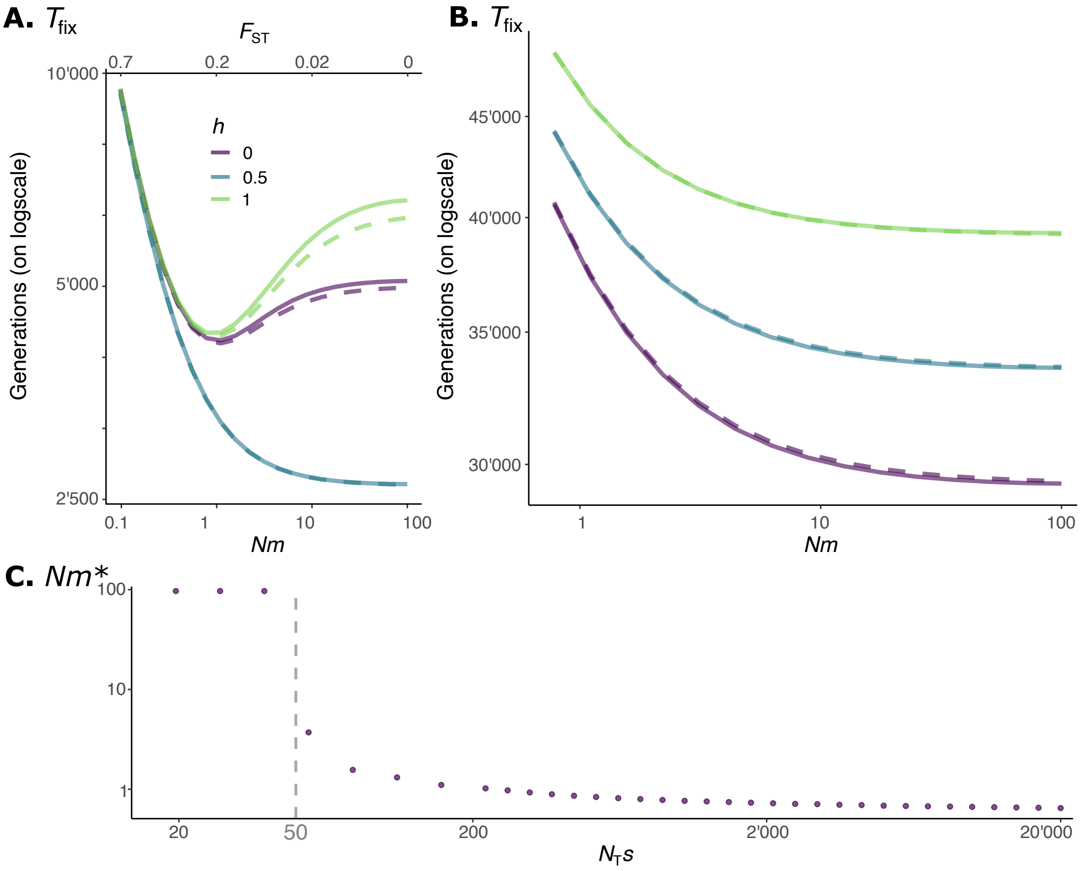


**Figure A: Selection threshold for faster fixation under dispersal limitation**. **A.** Expected time to fixation $T_{\text{fix}}(p_0)$ (on a log scale) of recessive ($h = 0$), additive ($h = 0.5$) and dominant ($h = 1$) alleles (in different colors, see legend) arising as single copies $p_0 = 1/(2N_T)$, with solid lines for general deme size and dispersal (i.e. using second column of Table 1) and dashed lines for large deme - weak dispersal limit (i.e. using third column of Table 1). Parameters: same as Figure 1, in particular $N_T s = 200$. **B.** Same as A, but with weaker selection, $N_T s = 2$ (so that $N_T s$ is as in Whitlock 2003). This shows that when selection is weak, fixation is always slower under limited dispersal, irrespective of dominance. **C.** Dispersal rate $m^*$ that minimizes the time to fixation $T_{\text{fix}}(p_0)$ of a recessive beneficial *de novo* mutation, according to the scaled selection coefficient $N_T s$. This was calculated based on numerical integration of eq. (7) for $N_T s < 200$. To speed up calculation, we used semi-deterministic approximation eq. (9) for $N_T s > 200$. This graph shows that when $N_T s$ is small, $Nm^* = 100$, i.e. fixation is fastest under panmixia and limited dispersal increases the time to fixation. But provided $N_T s > 50$ approximately (vertical dashed line), $Nm^* < 100$, indicating that limited dispersal reduces the time to fixation. Other parameters: same as Figure 1.

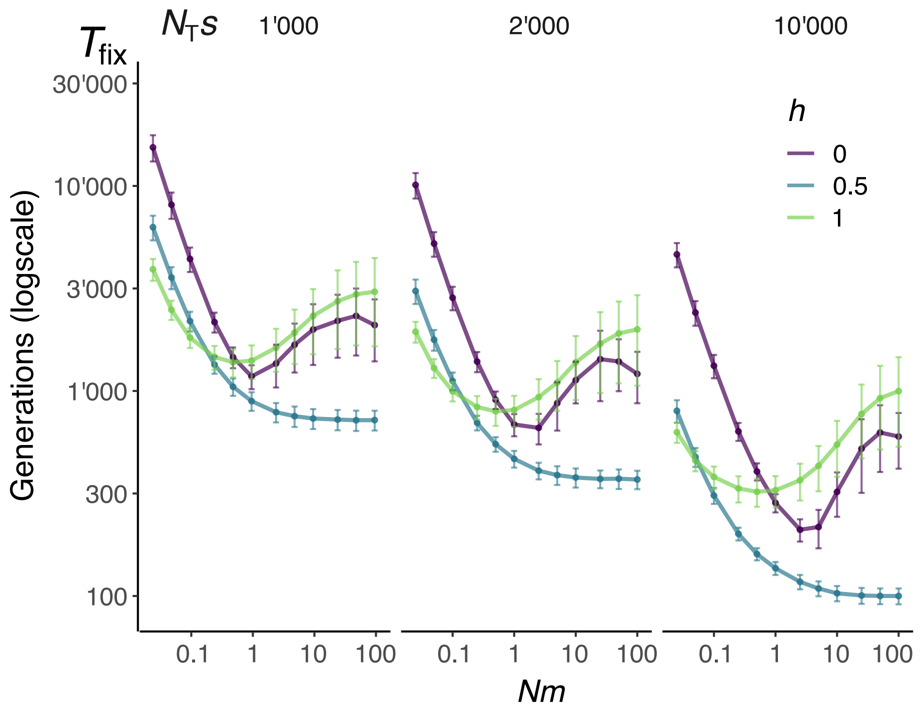


**Figure B: Fixation time under strong selection.** Average time to fixation (on a log scale) of recessive ($h = 0$), additive ($h = 0.5$), and dominant ($h = 1$) alleles arising as single copies $p_0 = 1/(2N_T)$, for various fecundity advantage $s$ (columns) from simulations (fixation events among at least 40'000 replicates for each set of parameters, error bars show standard deviation, section B for details). Parameters: $N_d = 200$, $N = 100$. As expected, stronger selection decreases time to fixation, but for each $N_T s$ investigated here, mild dispersal limitation speeds up the fixation of non-additive alleles (as in our baseline model).

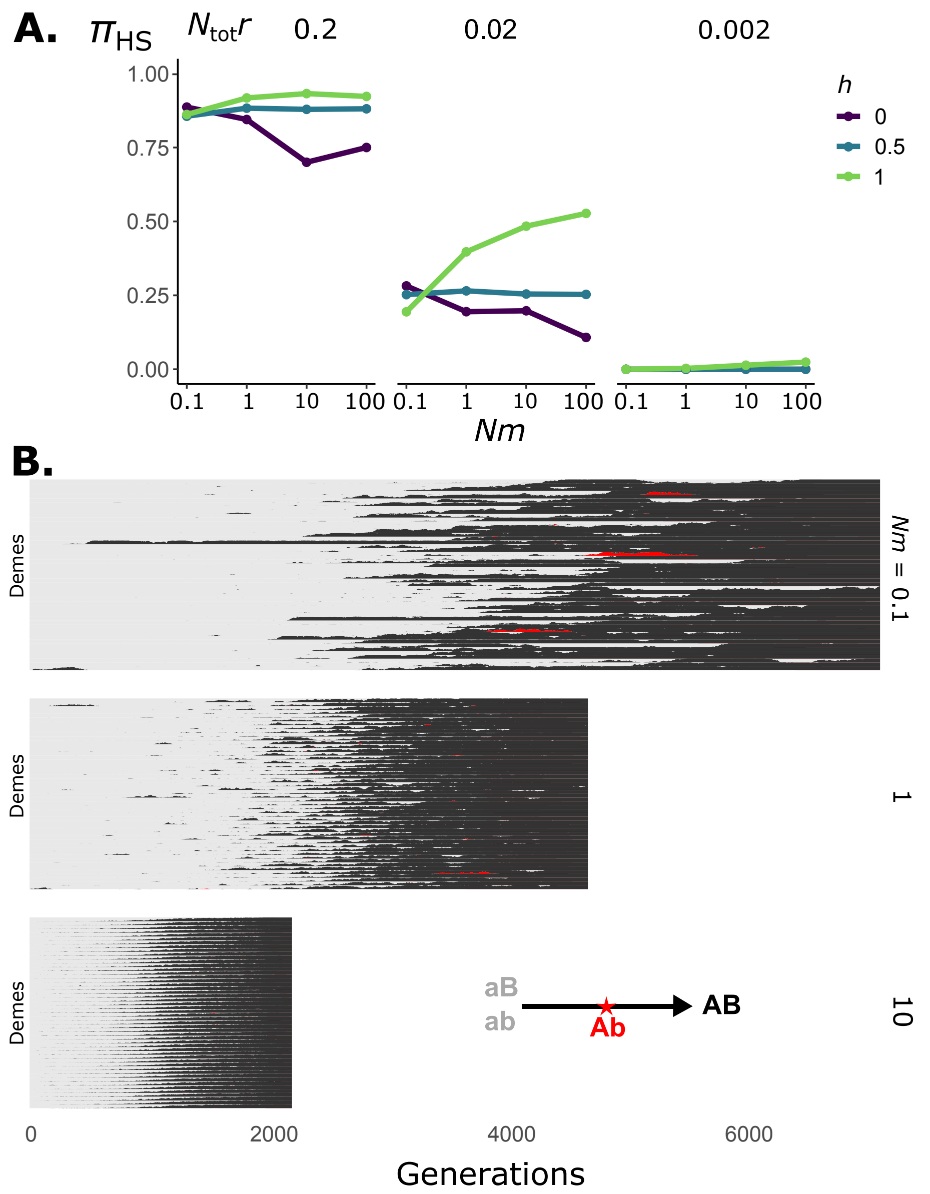


**Figure C: Effects of limited dispersal on the probability of hard vs. soft sweeps**. **A.** Probability $\pi_{\text{HS}}$ of a hard sweep, i.e. that polymorphism at a neutral locus is lost with fixation of a beneficial allele at a linked locus (section D for details on simulations). Each column corresponds to a different rate of recombination between the selected and the neutral loci, see legend for level of dominance of beneficial allele. These graphs show that limited dispersal does not influence $\pi_{\text{HS}}$ under additive fitness effects ($h = 1/2$), but increases (decreases) $\pi_{\text{HS}}$ for recessive (dominant) effects. Other parameters: same as Figure 1. **B.** Evolutionary dynamics of hard sweeps in populations under additive effects ($h = 0.5$) for three different dispersal rates ($Nm = 0.1, 1, 10$). For each dispersal rate, each line corresponds to one deme (50 shown out of 200 for illustration), showing haplotype frequencies within that deme through time. The population is initially composed of maladapted haplotypes ($aB$ and $ab$ in gray) and eventually fixes haplotype $AB$ (in black) leading to a hard sweep. In red is shown the frequency of the alternative $Ab$ haplotype that is created through recombination. These plots show that although this $Ab$ haplotype emerges multiple times under low dispersal rates ($Nm = 0.1$ and $Nm = 1$), it is eventually lost due to genetic drift within demes. When dispersal rate is high ($Nm = 10$), there is not enough time for the $Ab$ haplotype to be generated by recombination. Parameters: $N_{\text{T}}r = 0.02$, other parameters: same as A.

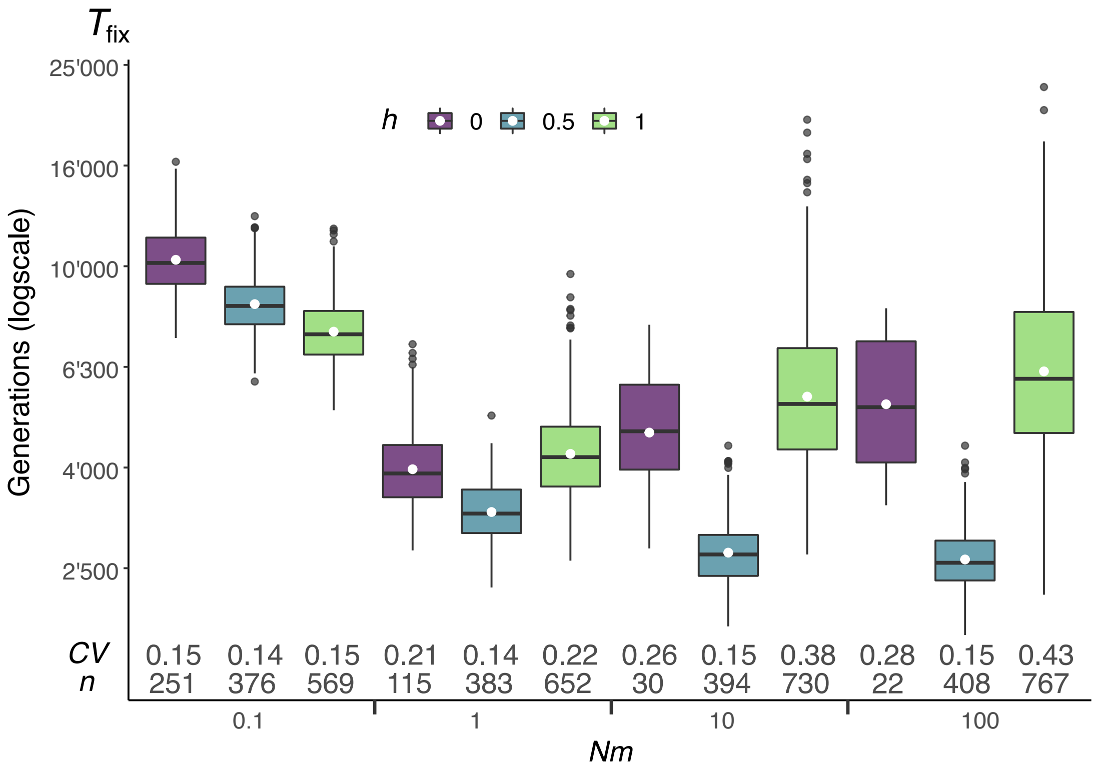


**Figure D: Summary statistics of fixation events**. Boxplot of the distribution of fixation times (given fixation) of a recessive ($h = 0$), additive ($h = 0.5$), and dominant ($h = 1$) allele for $Nm = 0.1, 1, 10$ and 100 (columns) for fixations among 40'000 replicates for each set of parameters. Boxes are from first quantile (Q1) to the third quantile (Q3). The median value (Q2) is shown with black line, and a white dot shows the mean. Outliers are shown as black dots. The coefficient of variation ($CV$) and the number of fixation events ($n$) are given for each set of parameters at the bottom of each boxplot. Parameters: same as Figure 1. Overall, these distributions are close to being symmetric around the mean.
